# Role Of Pcdh15 In The Development Of Intrinsic Polarity Of Inner Ear Hair Cells

**DOI:** 10.1101/2024.12.04.626923

**Authors:** Raman Kaushik, Shivangi Pandey, Anubhav Prakash, Takaya Abe, Hiroshi Kiyonari, Raj K. Ladher

**Affiliations:** National Centre for Biological Sciences, Tata Institute of Fundamental Research, GKVK PO, Bellary Road, Bangalore, India, 560065; Trivedi School of Biosciences, Ashoka University, Plot No. 2, Rajiv Gandhi Education City, National Capital Region P.O. Rai, Sonipat Haryana-131029; Laboratory for Animal Resources and Genetic Engineering, RIKEN Center for Biosystems Dynamics Research, 2-2-3 Minatojima-minamimachi, Chuo-ku, Kobe, Hyogo 6500047, Japan

**Keywords:** Inner Ear Hair Cell, Polarity, Protocadherin-15, Cochlea

## Abstract

In vertebrates, auditory information is transduced in the cochlea by mechanosensory hair cells (HC) through an eccentrically organised structure known as the hair bundle. This consists of a true cilium, known as the kinocilium, and modified microvilli, known as stereocilia. The hair bundle has a distinct structure with stereocilia organised in graded rows, with the longest abutting the kinocilia. The hair bundles of all HC are aligned to the tissue axis and are planar polarised. Important in the development and physiology of HC are protein bridges consisting of cadherin-23 and protocadherin-15. These link the tips of stereocilia, where they play a role in mechanotransduction, and between the kinocilia and the stereocilia, where they are involved in development. Both cadherin-23 and protocadherin-15 mutations result in defects in planar polarity; however, the mechanism through which this defect arises is unclear. Using a novel mutant for the Pcdh15-CD2 isoform, we show that while the initial deflection of the kinocilium occurs, its peripheral migration to register with Gαi3 is perturbed. Pcdh15-CD2 genetically interacts with Gpsm2, perturbing vestibular function. We find that the earliest expression of Pcdh15-CD2 is at the base of the kinocilia, and the defects in morphogenesis occur before the formation of kinocilial links. By re-introducing functional Pcdh15-CD2, we show that polarity can be restored, suggesting that Pcdh15-CD2 couples kinocilial and stereocilial development through a mechanism independent of kinocilial links.

## INTRODUCTION

Hearing has conferred numerous evolutionary advantages to animals, such as prey detection, predator avoidance and communication. Hearing perception occurs in the cochlea of the inner ear, in a specialised sensory epithelium known as the organ of Corti (OC). OC comprises four rows of mechanosensory hair cells (HC), divided into three rows of outer (OHC) and one row of inner hair cells (IHC), all surrounded by supporting cells [1, 2]. HC have hair bundles on their apical surface, which consist of graded rows of actin-based stereocilia increasing in height, with the tallest attached to a single true cilium, the kinocilium [3]. The development and orientation of the hair bundles are critical for the functioning of the inner ear as they are only mechanosensitive along one axis, the short-tall axis defined by the kinocilium [4]. Hair bundle development starts with centrally positioned kinocilium surrounded by microvilli covering the apical surface [5–8]. The kinocilium then centrifugally repositions to the periphery of the hair cell, followed by a re-orientation along the edge towards the abneural edge of the hair cells [9, 10]. Microvilli near the kinocilium then elongate and form the stereocilia, and the whole hair bundle moves towards the centre of the apical surface [11]. These initial steps of hair cell development define its intrinsic polarity. This intrinsic polarity is aligned across the OC such that the intrinsic polarity of all HC is aligned to the tissue axis, a process called planar cell polarity (PCP) [12].

Mutation studies have revealed a complex interplay of genes and pathways governing these processes. PCP is controlled by a molecularly conserved pathway of genes. In the cochlea, these comprise at least six different core PCP proteins [11, 12]. These proteins are expressed asymmetrically at the intercellular junctions and are crucial for the tissue-wide arrangement of hair cells. Intrinsic polarity of HC relies on the interaction of the kinocilium and a signalling complex initially defined in spindle positioning during asymmetric cell division. These proteins include the inhibitory G protein subunit alpha (Gαi), its binding partner GPSM2 (G protein signalling modulator) and Inscuteable (Insc) [13]. Mutation of Gαi, Gpsm2 or Insc in the inner ear only infrequently affects intrinsic polarity, although PCP is perturbed, and frequently the hair bundle is fragmented [9, 10, 14]. Similar defects have also been observed in mutants of genes involved in various aspects of cilia function, such as mutants of intraflagellar transport complex components or of molecules involved in kinocilia assembly [15–19]. Perturbations in the localisation of Gαi protein in the kinocilia mutants, Mkss, Bbs8 or Ift88 suggest a relationship between the kinocilia and the signalling mechanisms that are thought to underlie apical patterning of HC [10, 14, 17].

Protein links between its components ensure the cohesion of the hair bundle [20]. These include links between the stereocilia and between the kinocilia and stereocilia. Studies on the components of the kinocilial links, Cdh23 and Pcdh15 suggest a function of these molecules in the generation of HC polarity [21, 22]. Indeed, mutants of the CD2 isoform of Pcdh15 show planar polarity defects and the kinocilium is now distant from the tallest stereocilia [23]. In addition, blocking Fgf signalling, and thus preventing Pcdh15 from entering the kinocilium, also results in kinocilia-stereocilia decoupling [24]. However, kinocilial links only appear after the radial relocation of the kinocilia [25]. This may suggest a role for Pcdh15, independent of its function in kinocilial links. In this study, we generate a mouse where Pcdh15-CD2 is fused to an epitope tag. We find that Pcdh15-CD2 protein is initially localised around the base of the cilium prior to the peripheral relocation. Mutation of Pcdh15-CD2 alters the localisation of the Gαi, similar to kinocilial mutants. We find evidence of a genetic interaction with Gpsm2, and in mice mutant for both Gpsm2 and Pcdh15-CD2 we not only exaggerate polarity defects and bundle fragmentation, we now find defects in vestibular function. By re-expressing Pcdh15-CD2 we find that it is possible to rescue these defects up to 2 days after peripheral relocation. Our data suggests a previously unappreciated role for Pcdh15-CD2 during kinociliogenesis in coupling the kinocilium with Gpsm2/Gαi signalling.

## METHODS

### Animal use and care

Experiments on animals were performed according to Institutional Animal Ethics Committee guidelines. Pcdh15-n38 mice (Accession No.: CDB0186E: https://large.riken.jp/distribution/mutant-list.html) were generated by CRISPR/Cas9-mediated knock-in in zygotes as previously described [26]. Briefly, the following guide RNAs (gRNAs) were designed by using CRISPRdirect [27] to insert the knock-in construct: 5’ gRNA target sequence – 5’-CTATATGCATAGCTTGCGTG-3’ and 3’ gRNA target sequence - 5’-CTCAGCAAGCACGTCTACGT. Founder mice were bred with C57BL/6N mice to generate heterozygous animals. Pcdh15-n38YF mice were made by crossing homozygous Pcdh15n38 with Pgk1-Cre ([28]: JAX#020811) mice. GPSM2 mice (Accession No. CDB1144K) were received from RIKEN BDR, and were the kind gift of Fumio Matsuzaki [29]. Atoh1-Cre/ESR1 (JAX#007684) mice were obtained from JAX [30]. All primers used for genotyping are detailed in Table S1.

### Immunofluorescence

Inner ear dissections were performed as previously described [31, 32]. Fixation was based on the choice of antibody use (Table S2). Fixed inner ears were washed with 1X PBS and further dissected to expose the sensory epithelium. Tectorial membrane was removed before permeabilisation with PBS + 0.3%Tween-20. Tissues were blocked with blocking buffer (10% goat serum, 1% bovine serum albumin (BSA) in permeabilisation buffer) and then incubated with primary antibodies (Table S2) overnight at 4°C. Tissues were then washed 5-6 times with PBST before incubating with secondary antibodies and phalloidin (Table S3) at room temperature for 1 hour. Tissues were washed 5-6 times before mounting with aqueous mounting media.

### Image acquisition and processing

Confocal images were captured by Olympus FV3000 inverted confocal microscope with a 60x objective (1.42 NA) using its software Olympus FV31S-SW. All images were captured at sub-saturation level under Nyquist sampling criteria. High-resolution images were taken on a Zeiss LSM 980 Airyscan 2 microscope with a 63x objective (1.4 NA). All the raw files were processed by ImageJ software.

### Immunoprecipitation and western blotting

For immunoprecipitation (IP), 20 mice cochlea at P5 stage were dissected and homogenised in lysis buffer (0.5% Triton X-100, 150mM NaCl, 5mM EDTA and 1X MS-SAFE protease and phosphatase inhibitor in PBS). Lysates were cleared by centrifugation at 15,000 g for 10 mins. Anti-HA or normal mouse IgG antibody was incubated with Dynabead^TM^ Protein G beads (Invitrogen) for 1 hr at 4°C to form the Dynabead-bound antibody complexes. Parallelly, the lysate was incubated with Dynabeads for reducing the non-specific binding. The cleared lysates were incubated with Dynabead bound antibody complexes of either HA-specific antibody (Sigma H3663) or with normal mouse IgG overnight at 4°C. The proteins were purified and washed 5 times with lysis buffer before eluting in 1X SDS loading buffer. Proteins were resolved by 10% SDS-PAGE and transferred on to PVDF membranes. The membranes were blocked by 5% BSA in 0.1% TBST buffer and incubated with primary antibodies overnight at 4°C. After 5 washes with 0.1% TBST buffer, membranes were incubated with peroxidase-conjugated secondary antibodies. Immunoreactive proteins were detected with western bright ECL detection reagents.

### Vestibular test

Littermate mice (5 months old) for different genotypes (Pcdh15^n38/+^:: Gpsm2^+/-^, Pcdh15^n38/n38^:: Gpsm2^+/-^ & Pcdh15^n38/+^:: Gpsm2^-/-^ & Pcdh15^n38/n38^:: Gpsm2^-/-^) were kept in a controlled environment. Vestibular test experiments were performed in a box (22cm X 15cm X 13cm) and the activity of mice was recorded for 5 min. The total distance covered, mean speed and no. of rotations were measured. Analyses were performed using ANY-maze software.

### Image analysis and statistics

Data were collected from at least three animals (from at least two different breeder pairs) for each experiment. Images were analysed using ImageJ.

For hair bundle defects, kinocilia position (marked with Arl13b or acetylated tubulin antibodies) was measured with respect to the hair cell cortex (marked by phalloidin). A vector was drawn from the centre of the hair cell to the kinocilium base whose length (r) is normalised with cell radius and the angle (θ) from the proximal to distal (P-D) axis. The kinocilia position in different hair cells was visualised by using polar plots in Matplotlib library of Python. Similarly, the frequencies of kinocilia orientation were measured and plotted from 0°-360° within 10° brackets.

For alpha – theta (α - θ) correlation analysis, θ angles were plotted against α angles. θ angles were calculated for angle of the kinocilia base from 0° (along the P-D axis), and α angles were calculated for the centre of the Gαi expression domains from the horizontal (0°) on the P-D axis.

Box and whisker graphs were plotted using GraphPad Prism 9.5.0. The same software was used for statistical analyses. We analysed data using ordinary one-way or two-way ANOVA test (Tukey’s multiple comparison test). P values were represented in the graphs; p <0.05 was considered statistically significant.

### Scanning electron microscopy (SEM)

Samples for SEM were prepared as described previously [32]. After inner ear dissection in 1X PBS, tissues were fixed in 2.5% glutaraldehyde in sodium cacodylate buffer (0.1M) with calcium chloride (3mM) for 48 hours. After fixation, samples were micro-dissected to expose the sensory epithelium. Samples were again processed with OTOTO method for an alternate series of fixation with 1% osmium tetroxide (O) and 0.5% thiocarbohydrazide (T). Samples were dehydrated first with serially increased alcohol percentage followed by critical point dehydration (Leica EM CPD300). Mounting was done on double-sided carbon adhesive tapes, followed by sputter coating. Samples were imaged on Zeiss Merlin Compact VP with SE2 detectors.

## RESULTS

### Generation of Pcdh15^n38/n38^ and Pcdh15^n38YF/n38YF^ mice

Our previous work had suggested that tyrosine phosphorylation on the cytoplasmic domain of the PCDH15-CD2 isoform was important for Pcdh15 function [24]. We hypothesised that finding interactions specific to phosphorylation to Pcdh15 would help to understand this function. To do this, we engineered a mouse: the endogenous exon 38 was replaced by a construct that consisted of a CD2 domain with an in-frame FLAG tag. In addition, the construct contained a mutant CD2 in which all 6 tyrosine residues were changed to phenylalanine, and fused to a HA tag. These sequences were flanked by Lox P, Lox71 and Lox 66 sites. Without Cre recombination, the wild-type Pcdh15-CD2 tagged with the FLAG epitope should be expressed (Pcdh15^n38/n38^) (Fig. 1A, S1A).

**Figure 1.**
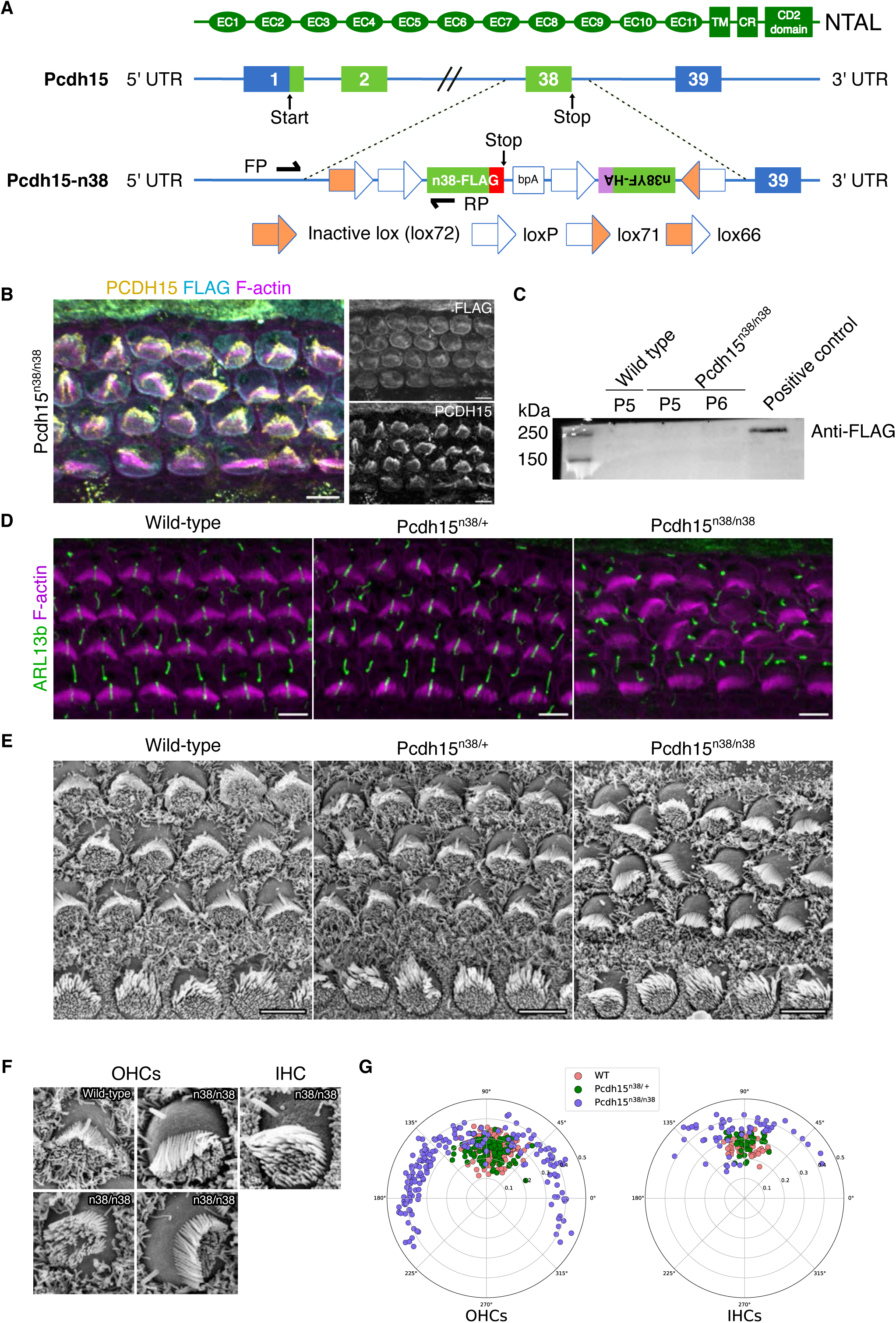
Pcdh15^n38/n38^ mice are functionally null and act similar to Pcdh15-ΔCD2 mutants. (A) Schematic of the DNA construct used to generate Pcdh15^n38/n38^ mice. (B) The organ of Corti of Pcdh15^n38/n38^ mice at P0 (mid). Immunostaining fails to detect the PCDH15-CD2-FLAG signal (cyan), although endogenous PCDH15 (yellow) is localised at the tip of stereocilia (magenta, marked with phalloidin). (C) Western blot fails to detect PCDH15-CD2-FLAG protein in Pcdh15^n38/n38^ mice OC (P5/6). Positive control show HEK293T cells lysate transfected with PCDH15-CD2-FLAG cDNA. (D-E) Fluorescence (D) and SEM (E) imaging shows hair bundle polarity defects in Pcdh15^n38/n38^ OC when compared to normal hair bundle orientation in WT & Pcdh15^n38/+^ (P0, mid). Cilia is marked with ARL13b (green) and stereocilia with phalloidin (magenta). (F) Kinocilium is dissociated from stereocilia in OHC and IHC of Pcdh15^n38/n38^ OC when compared to WT. (G) Polar plots of kinocilia positions in hair cells show higher circular standard deviation, indicative of hair bundle polarity defects in Pcdh15^n38/n38^ OC (slate blue). This is much reduced in WT (pink) & Pcdh15^n38/+^ (green) (P0, mid). (Scale bar = 5 μm).

To check for PCDH15-CD2 expression in hair cells of the organ of Corti, we performed immuno-localisation with a FLAG-tag specific antibody in Pcdh15^n38/n38^ mice. While PCDH15 expression could be visualised in stereocilia tips, we were unable to detect immuno-reactivity for the FLAG-tag (Fig. 1B). The lack of FLAG immunoreactivity in the tissue was confirmed by Western blotting. We failed to detect the FLAG epitope in P5/6 Pcdh15^n38/n38^ organ of Corti lysates, despite detection in HEK cells (Fig. 1C, S1B). RT-PCR indicated no difference in mRNA expression between WT and Pcdh15^n38/n38^ mice (Fig. S1C). This suggested that translation of the mutant Pcdh15-CD2 isoform was impaired, indicating that the Pcdh15^n38/n38^ mice are functionally null and act similar to Pcdh15-ΔCD2 mutants reported earlier [23, 33].

### Defects in hair bundle polarity in Pcdh15^n38/n38^ mice

Pcdh15-ΔCD2 mutants show defects in hair bundle orientation and mechanotransduction [23, 33]. To ask if Pcdh15^n38/n38^ mice show a similar phenotype to Pcdh15-ΔCD2, we immunostained the P0 organ of Corti with Arl13b and phalloidin to determine hair bundle polarity. The average kinocilium deviation from the PCP axis was comparable in P0 wild-type and Pcdh15^n38/+^ animals (Fig. 1D-E & 1G). In OHCs this deviation, at P0, is 13° ± 11° in Pcdh15^n38/+^; however, in Pcdh15^n38/n38^ animals, this deviation is 59° ± 32° indicating a defect in planar polarity. While hair bundle polarity is affected equally in all three rows of OHCs, IHCs show a less severe defect in hair bundle polarity, with the average kinocilium deviation of 23° ± 15° in Pcdh15^n38/n38^ as compared to 8° ± 6° in Pcdh15^n38/+^ animals (Fig. 1G). Despite the defect in polarity, the expression of the core PCP protein, Vangl2 is unchanged in Pcdh15^n38/n38^ mice (Fig. S1D). Closer inspection using SEM revealed a decoupling of the kinocilium from the stereocilia (Fig. 1E). In Pcdh15^n38/+^, almost all kinocilia are attached to the longest stereocilia and are found at the vertex of the stereocilia bundle. In Pcdh15^n38/n38^ mice, we were unable to detect any hair cell with kinocilium coupled to the stereocilia. Additionally, we observed misoriented stereocilia bundles of varying degrees from PCP axis, as well as occasional circular bundles (Fig. 1F).

### Tyrosine phosphorylation of the CD2 domain is not essential for Pcdh15 function in mice

Our previous work had identified that Fgfr1 signalling was involved in the coupling of the kinocilium to the stereocilia, through the phosphorylation of the cytoplasmic domain of Pcdh15-CD2 [24]. By mating Pcdh15^n38/n38^ with Pgk1-Cre, we generated Pcdh15^n38YF/+^ heterozygous animals, which were back-crossed to give homozygotes (Fig. 2A-B, S2A). These mice showed the epitope tag (HA) expression in both kinocilia and at stereocilia tips, indicating normal expression of the modified Pcdh15-CD2-HA (Fig. 2C-D). Although we could not resolve the hair bundle at embryonic stages, at postnatal stages, we observed PCDH15-CD2-HA expression at the tip of all three rows of stereocilia in HCs (Fig. S2B). This data was verified by immunoprecipitation from the cochlear lysate of Pcdh15^n38YF/n38YF^ mice, showing a specific band when compared to the IgG control (Fig. 2E, S2C). Arl13b and phalloidin staining was used to measure hair bundle polarity, together with SEM for determining kinocilia-stereocilia coupling. Despite Pcdh15^n38/n38^ showing defects in polarity, Pcdh15^n38YF/n38YF^ were similar to wild-type controls (Fig. 2F-H, S2D). This suggests that the phosphorylation of tyrosine in the CD2 domain of Pcdh15^n38YF/n38YF^ is not essential for normal Pcdh15 function.

**Figure 2.**
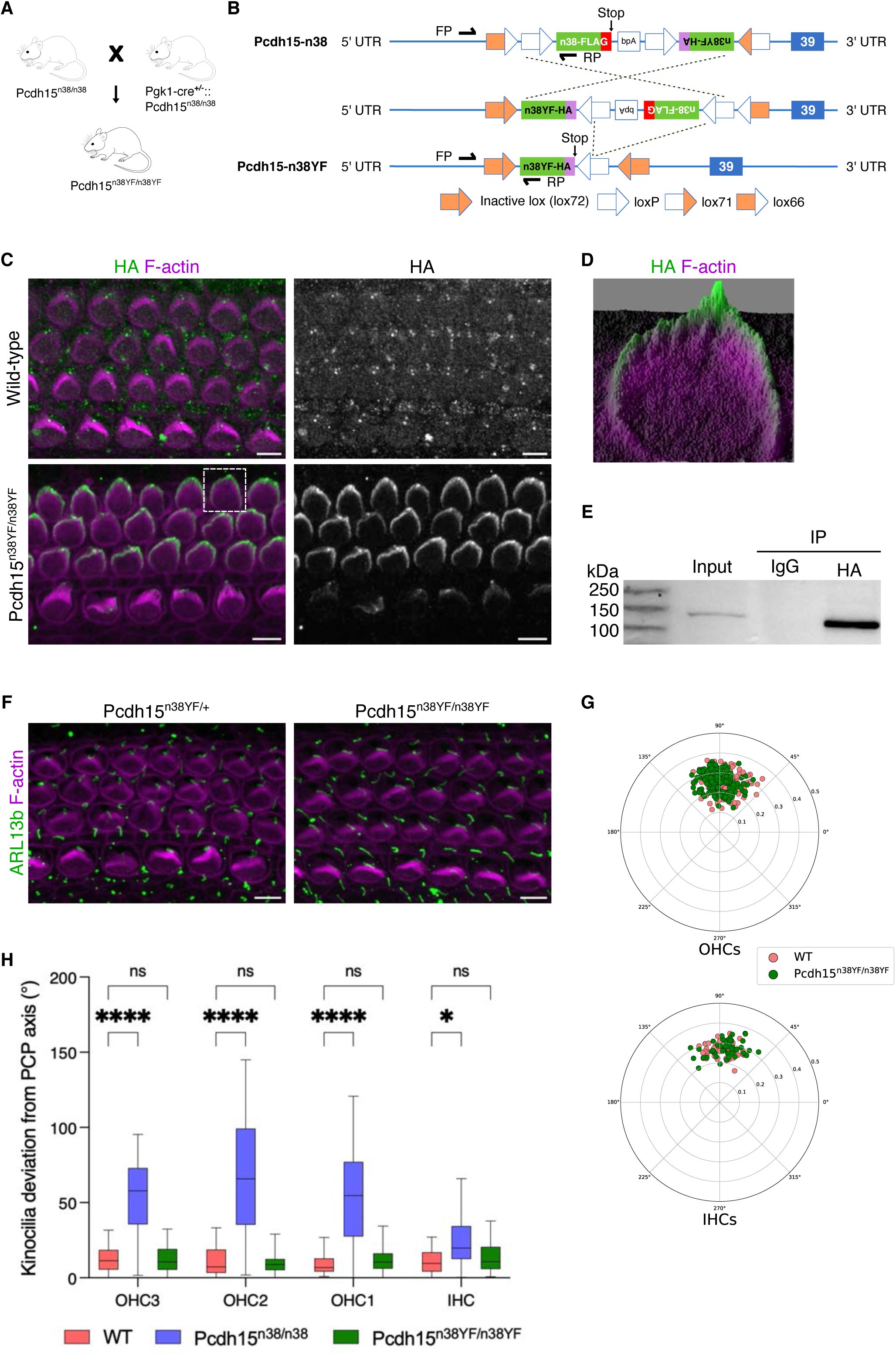
Tyrosine phosphorylation of the CD2 domain is not essential for PCDH15 function. (A) Schematic of the mice breeding scheme to produce Pcdh15^n38YF/n38YF^ mice. (B) Schematic of the Pcdh15-n38 DNA construct recombination to Pcdh15-n38YF under Cre activity. (C) Modified PCDH15-CD2-HA (green) is expressed in the stereocilia (magenta) of Pcdh15^n38YF/n38YF^ hair cells (P0, mid), but not in WT. (D) Surface plot of OHC shows the PCDH15-CD2-HA (green) localisation at the tip of hair bundle. (E) IP assay shows that PCDH15-CD2-HA is precipitated with anti-HA antibody. (F) Hair bundle polarity is normal in Pcdh15^n38YF/n38YF^ and Pcdh15^n38YF/+^ hair cells (P0, mid). Cilia is marked with ARL13b (green) and stereocilia with phalloidin (magenta). (G) Polar plots marking kinocilia positions in hair cells show the normal hair bundle polarity in Pcdh15^n38YF/n38YF^ OC (green) (P0, mid). (H) Kinocilia are aligned normally with PCP axis in WT (pink) and Pcdh15^n38YF/n38YF^ (green) hair cells, but significantly deviated in Pcdh15^n38/n38^ (slate blue). The deviation is greater in OHCs when compared to IHCs (P0, mid). (Scale bar = 5 μm, Two-way Anova with Tukey’s multiple comparisons test, ns = P>0.05, * = P<0.05, ** = P<0.01, *** = P<0.001, **** = P<0.0001).

Pcdh15-CD2 has also been implicated in mechanotransduction, although the MET channels in Pcdh15-ΔCD2 are functional [23]. Using the styryl pyridinium dye, FM1-43 FX which stains hair cells after entering through MET channels, we asked if MET channels were functional [34]. We find that HC in the P5 organ of Corti from both Pcdh15^n38/n38^ and Pcdh15^n38YF/n38YF^ mice are stained by FM1-43, suggesting that MET channels can open (Fig. S2E). Using the Preyer’s reflex, we found that P60 Pcdh15^n38/n38^ mice could not respond to sound (data not shown). This data suggests that even though MET channels in Pcdh15^n38/n38^ mice can open, mutants cannot hear.

### PCDH15-CD2 is expressed in the kinocilium prior to its off-centre movement

To understand the role of Pcdh15-CD2 in intrinsic HC polarity, we utilised the HA tag on Pcdh15^n38YF/n38YF^ to map its earliest expression during hair bundle morphogenesis. At E15.5, we observe PCDH15-CD2-HA expression at the base of the organ of Corti, in the IHC kinocilium which at these stages has yet to move off-centre (Fig. 3A). We also noted HA positive cortical punctae around the kinocilia (Fig. 3A). At these stages we did not observe HA expression in the OHC or any HC at the apex or mid of the organ of Corti. At E16.5, HA-staining could be observed in the kinocilia of both IHC and OHC in apex of organ of Corti (Fig. 3B).

**Figure 3.**
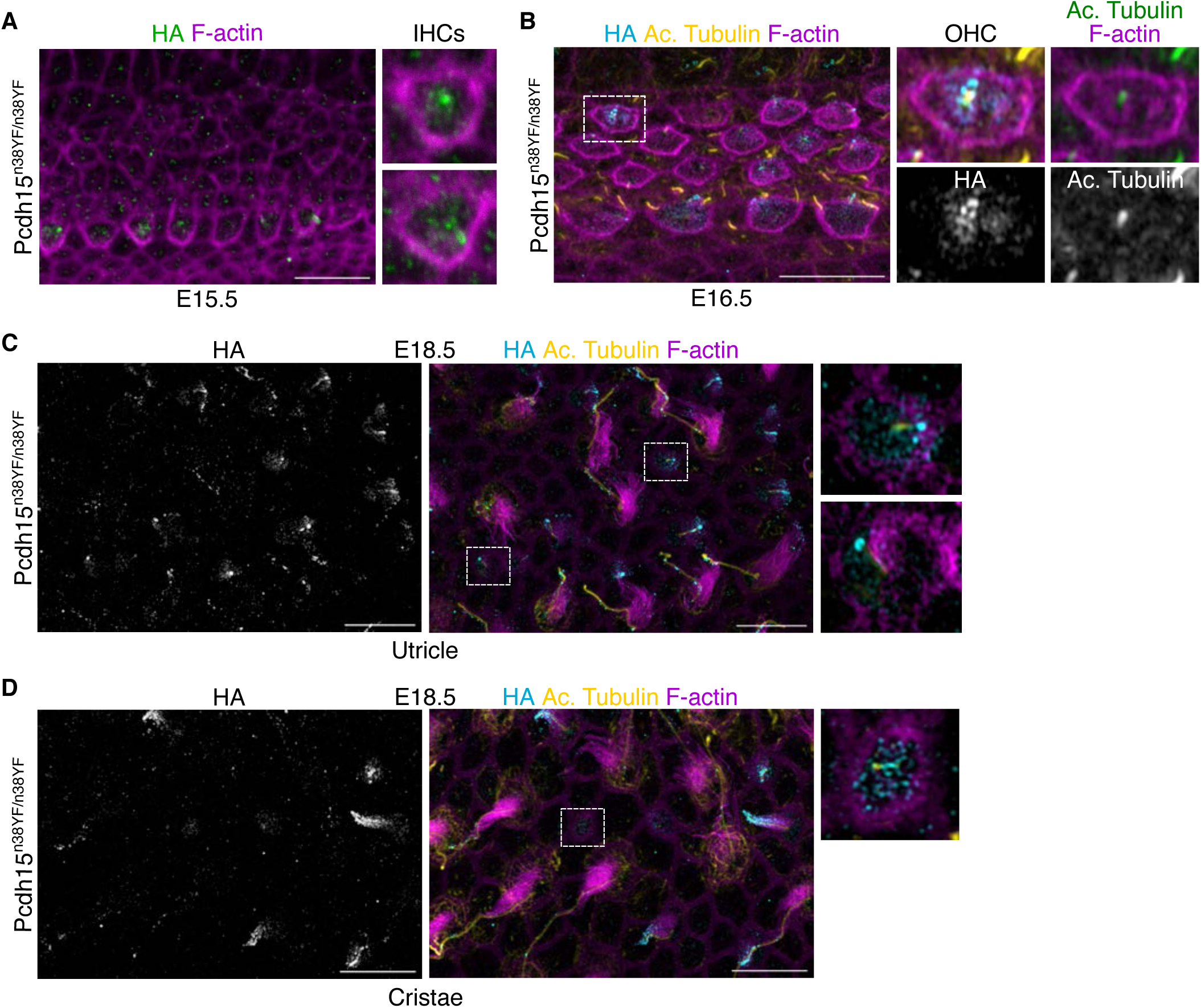
PCDH15-CD2-HA is expressed in the kinocilium prior to its off-centre movement. (A) PCDH15-CD2-HA (green) is localised in centrally located kinocilium of IHCs, but not expressed in OHCs of E15.5 OC (base). (B) PCDH15-CD2-HA (cyan) is localised in centrally located kinocilium (yellow, marked with acetylated tubulin) of OHCs of E16.5 OC (apex). (C - D) PCDH15-CD2-HA (cyan) is localised in centrally located kinocilium (yellow, marked with acetylated tubulin) of newly differentiating HCs of E18.5 utricle (C) and (D) cristae. As hair cell mature, this expression is enriched in hair bundles (magenta, marked with phalloidin) and becomes refined. (Scale bar = 10 μm).

To ask if PCDH15-CD2-HA was also expressed on the hair bundle of vestibular HC, we assessed the utricle and cristae. At E18.5 newly differentiating hair cells can be distinguished from mature HC. In newly differentiating HC, PCDH15-CD2-HA expression is found in the kinocilium, prior to its off-centre movement. Similar to the auditory HC, PCDH15-CD2-HA expression was observed in punctae around the kinocilium of HC of the utricle (Fig. 3C) and cristae at E18.5 (Fig. 3D). In more mature HC, distinct punctae in kinocilia are visible.

### Pcdh15^n38/n38^ mice exhibits early hair bundle polarity defects

During the development of the hair cell, the kinocilium initially moves to an eccentric location, adjusts along the periphery, and then relocalises towards the centroid of the HC apex with the expansion of the bare zone [11]. As PCDH15-CD2 expression could be detected in kinocilia before the off-centre movement, we asked at which point polarity defects arose. We determined kinocilium position using Arl13b at E16.5 and E18.5. In Pcdh15^n38/+^ at E16.5, and prior to its off-centre movement, kinocilium position is distributed around the centroid of the HC with a slight bias to the abneural side of the OC. In contrast, the kinocilium does not show such a bias in Pcdh15^n38/n38^ mice (Fig. 4A-C). We next assessed the kinocilia position pattern at E18.5. Pcdh15^n38/+^ kinocilia had relocalised towards the centroid of the HC. In contrast, the kinocilia of Pcdh15^n38/n38^ mice are distributed randomly and remain positioned near the periphery of hair cells (Fig. 4D-F). To further understand the randomised kinocilia position we investigate the position of centrioles. In Pcdh15^n38/+^ HCs at E18.5, centrioles of the basal body are aligned at a right angles to each other, with the mother centrioles attached to the kinocilia, and the daughter centriole extending basally. In contrast, in Pcdh15^n38/n38^ the centrioles are separated from each other in Pcdh15^n38/n38^ mice, and although the mother centriole is still in contact with kinocilia, the daughter centriole position is randomised (Fig. 4G-H). Taken together, these data suggest that although the initial off-centre movement occurs, subsequent relocalisation is impaired.

**Figure 4.**
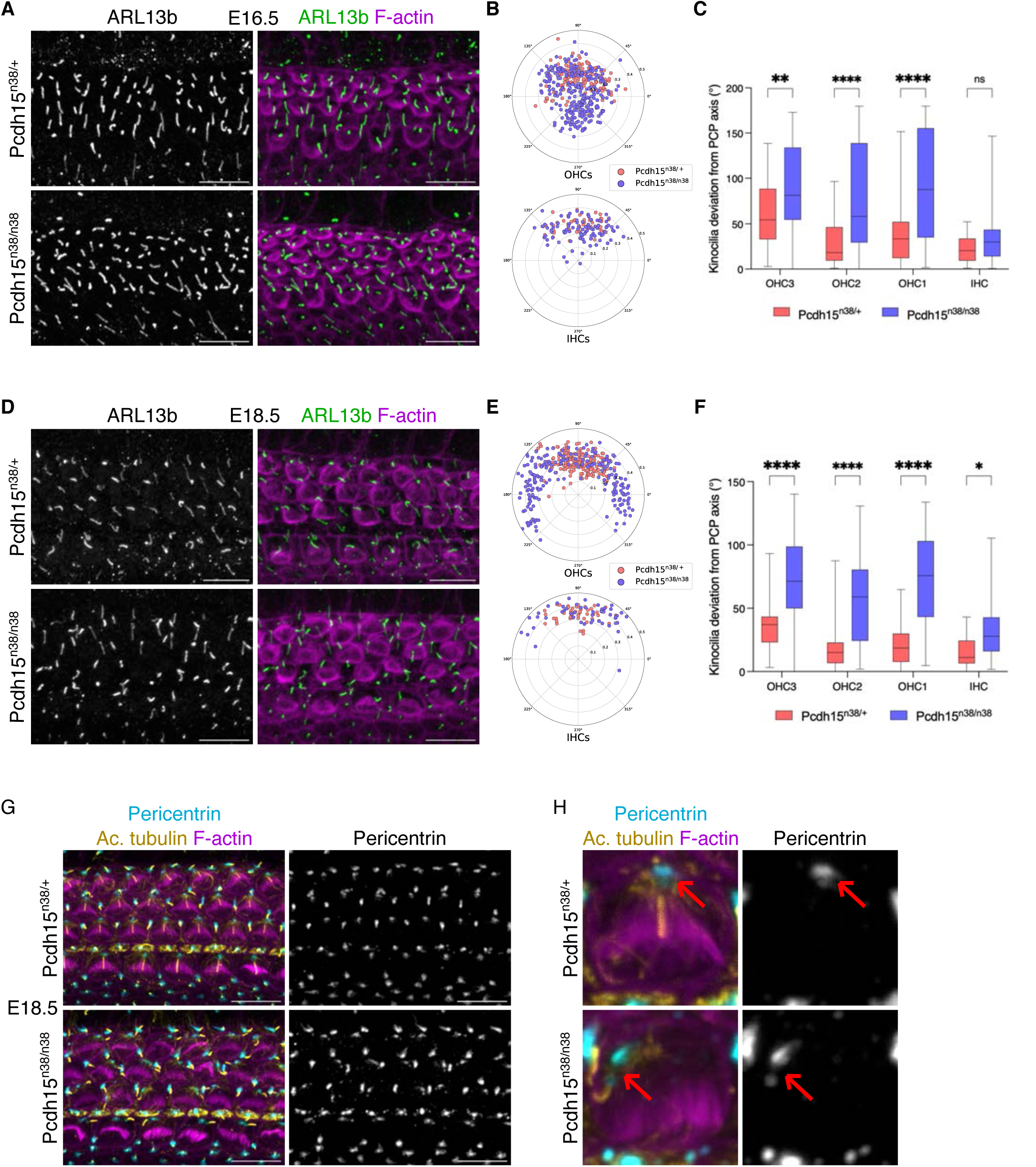
Pcdh15^n38/n38^ mice exhibits early hair bundle polarity defects. (A) Hair bundle polarity is perturbed in Pcdh15^n38/n38^ hair cells of E16.5 OC (mid). Cilia is marked with Arl13b (green) and stereocilia with phalloidin (magenta). (Scale bar = 10 μm). (B) Polar projections marking kinocilia positions in E16.5 stage (mid) hair cells show the perturbed hair bundle polarity in Pcdh15^n38/n38^ (slate blue). (C) Kinocilia are deviated significantly from the PCP axis in Pcdh15^n38/n38^ (slate blue) OHCs as compared to Pcdh15^n38/+^ (pink) (E16.5, mid). (Two-way Anova with Tukey’s multiple comparisons test, ns = P>0.05, * = P<0.05, ** = P<0.01, *** = P<0.001, **** = P<0.0001). (D) The perturbation in hair bundle polarity continues in Pcdh15^n38/n38^ hair cells of E18.5 OC (mid). Cilia is marked with Arl13b (green) and stereocilia with phalloidin (magenta). (Scale bar = 10 μm). (E) Kinocilia positions in E18.5 stage (mid) hair cells show the perturbed hair bundle polarity in Pcdh15^n38/n38^ (slate blue). (F) Kinocilia are deviated significantly from the PCP axis in Pcdh15^n38/n38^ (slate blue) HCs (E18.5, mid). (Two-way Anova with Tukey’s multiple comparisons test, ns = P>0.05, * = P<0.05, ** = P<0.01, *** = P<0.001, **** = P<0.0001). (G-H) The centrioles (cyan, marked with pericentrin) are not positioned at right angles to each other in HCs of Pcdh15^n38/n38^ mice (E18.5, base). (Scale bar = 10 μm).

### The coordination of kinocilium with Gαi expression is lost in Pcdh15-CD2 mutants

Intrinsic polarity is controlled through a pathway involving Gpsm2 and Gαi. Gαi expression is normally restricted to a narrow lateral crescent in HC [9, 10, 14]. The crescent not only marks intrinsic polarity, it also specifies the bare zone, a region free of microvilli, lateral to the kinocilium [9]. In mutants for the ciliary genes Bbs8, Ift88 or McKusick-Kaufman syndrome (Mkks) gene, the expression domain of Gαi is expanded, so that it covers the entire lateral half of the HC [10, 14, 17]. Similar to ciliary mutants, we find that the loss of Pcdh15-CD2 also expands Gαi expression medially (Fig. 5A-C).

**Figure 5.**
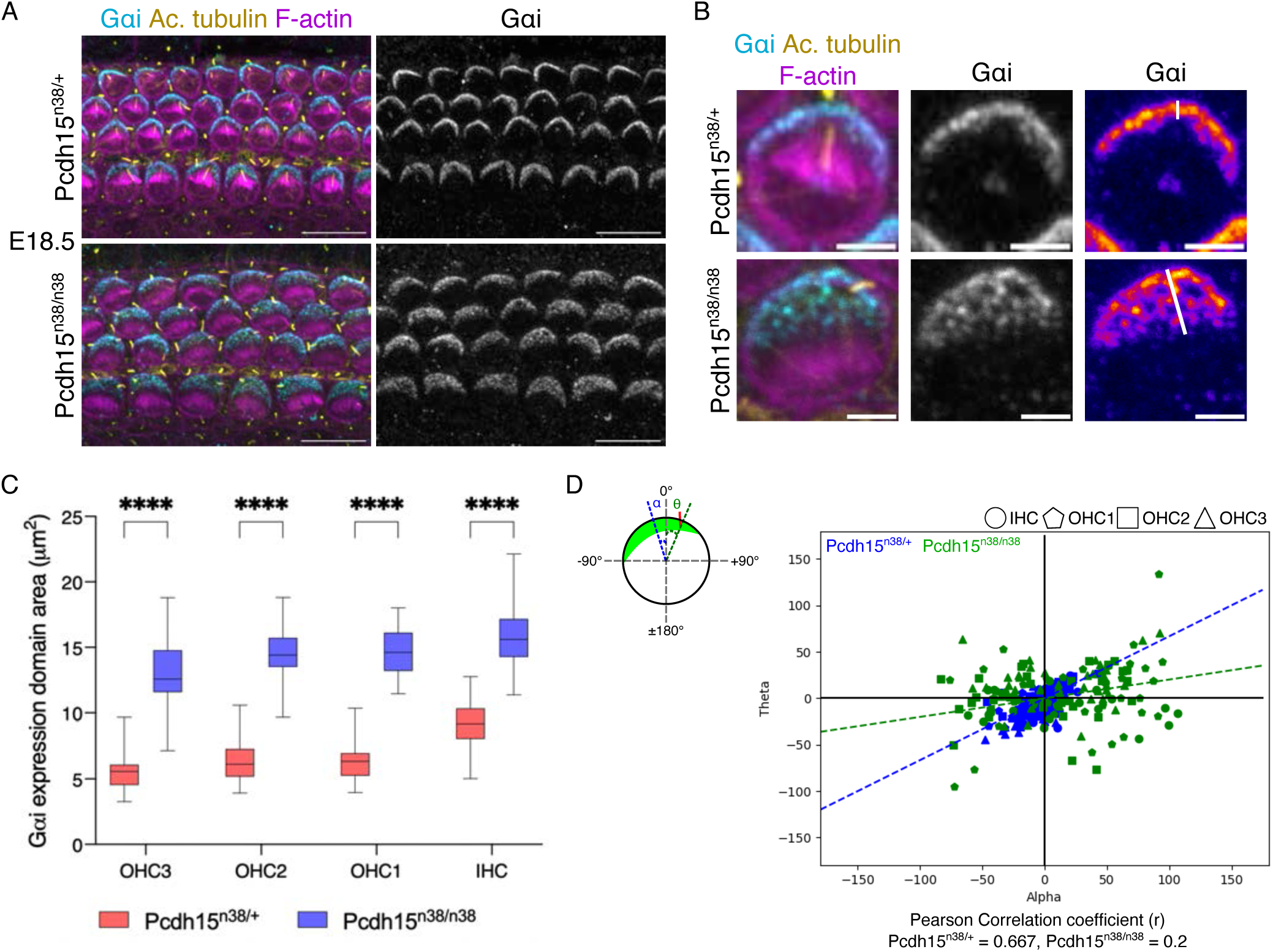
Kinocilium position is not coordinated with Gαi expression in Pcdh15^n38/n38^ mice hair cells. (A) Gαi expression (cyan) is perturbed with respect to the kinocilium position (yellow, marked with acetylated tubulin) in E18.5 (base) hair cells of Pcdh15^n38/n38^ mice. (Scale bar = 10 μm). (B) Gαi expression domain (cyan) is extended medially in HCs of Pcdh15^n38/n38^ mice. (Scale bar = 2 μm). (C) In comparison to Pcdh15^n38/+^ mice, Gαi expression domain is spread significantly in HCs of Pcdh15^n38/n38^ mice (E18.5, base). (Two-way Anova with Tukey’s multiple comparisons test, ns = P>0.05, * = P<0.05, ** = P<0.01, *** = P<0.001, **** = P<0.0001). (D) Correlation between the kinocilium position and Gαi expression domain is reduced in HCs of Pcdh15^n38/n38^ mice (green) when compared to Pcdh15^n38/+^ (blue) (E18.5, base). Alpha (α) is the angle measured for kinocilium with PCP axis and theta (θ) is the angle measured for Gαi expression domain (mid) with PCP axis.

After its off-centre movement, the kinocilium undergoes peripheral relocation, such that it is situated in the middle of a domain of Gαi expression. In Mkks mutants, this shift is perturbed [10]. To ask if a similar effect was visible in the Pcdh15-CD2 mutants, we measured the correlation between angle the kinocilia makes with the tissue axis (theta) and the angle that the midpoint of Gαi expression makes with the tissue axis (alpha) In Pcdh15^n38/+^, strong angular correlation between centre of the Gαi expression domain and kinocilia is observed, with a Pearson correlation coefficient of 0.667. In Pcdh15^n38/n38^ mutant mice this correlation coefficient is reduced to 0.2 (Fig. 5D), suggesting that Pcdh15^n38/n38^ mutant HC fail to undergo a peripheral relocation during hair bundle development, remaining decoupled from Gαi.

### Pcdh15-CD2 and GPSM2 proteins operate cooperatively in hair bundle development

We next asked if PCDH15-CD2 and Gαi-GPSM2 showed any genetic interaction. We used a null Gpsm2 mutant, in which almost the entire coding sequence is deleted [29]. On a Pcdh15^n38/+^ background, neither Gpsm2^+/-^ heterozygotes (10° ± 8°) nor Gpsm2^-/-^ homozygotes (7° ± 6°) show significant defects in planar polarity at P1 (Fig. 6A, C & E). Pcdh15^n38/n38^ mice show defects in hair bundle polarity (Fig. 1G: quant). However, both Pcdh15^n38/n38^:: Gpsm2^+/-^ and Pcdh15^n38/n38^:: Gpsm2^-/-^ compound mutants showed an exacerbated polarity defect (67° ± 43° and 59° ± 39° respectively) (Fig. 6 B, D, E). Gpsm2 mutants HC show hair bundle fragmentation (24% of OHCs & 14% of IHCs) (Fig. 6F) [10, 14]. We thus asked if the number of hair cells showing bundle fragmentation increased in the compound mutants. We found increased fragmentation in the compound mutants, with OHCs more affected than IHC (46% of OHCs & 28% IHCs) (Fig. 6F, S3). Moreover, and in contrast to planar polarity, the proportion of HCs in the Pcdh15^n38/n38^:: Gpsm2^-/-^ organ of Corti that shows bundle fragmentation is significantly higher than in Pcdh15^n38/n38^:: Gpsm2^+/-^ and Pcdh15^n38/+^:: Gpsm2^-/-^ cochlea (Fig. 6F). Individually, Gpsm2 null and Pcdh15^n38/n38^ mice do not exhibit vestibular defects, however, we observed Pcdh15^n38/n38^:: Gpsm2^-/-^ mice show circling behaviour and hyperactivity (Fig. 6G: Supp Move 1-4). In an open field test, we found that Pcdh15^n38/n38^:: Gpsm2^-/-^ mice travel larger distances, with higher mean speed and show a larger number of rotations, with a preference to anti-clockwise turns (Fig. 6H-J). Taken together, these data suggest a genetic interaction between Pcdh15 and Gpsm2.

**Figure 6.**
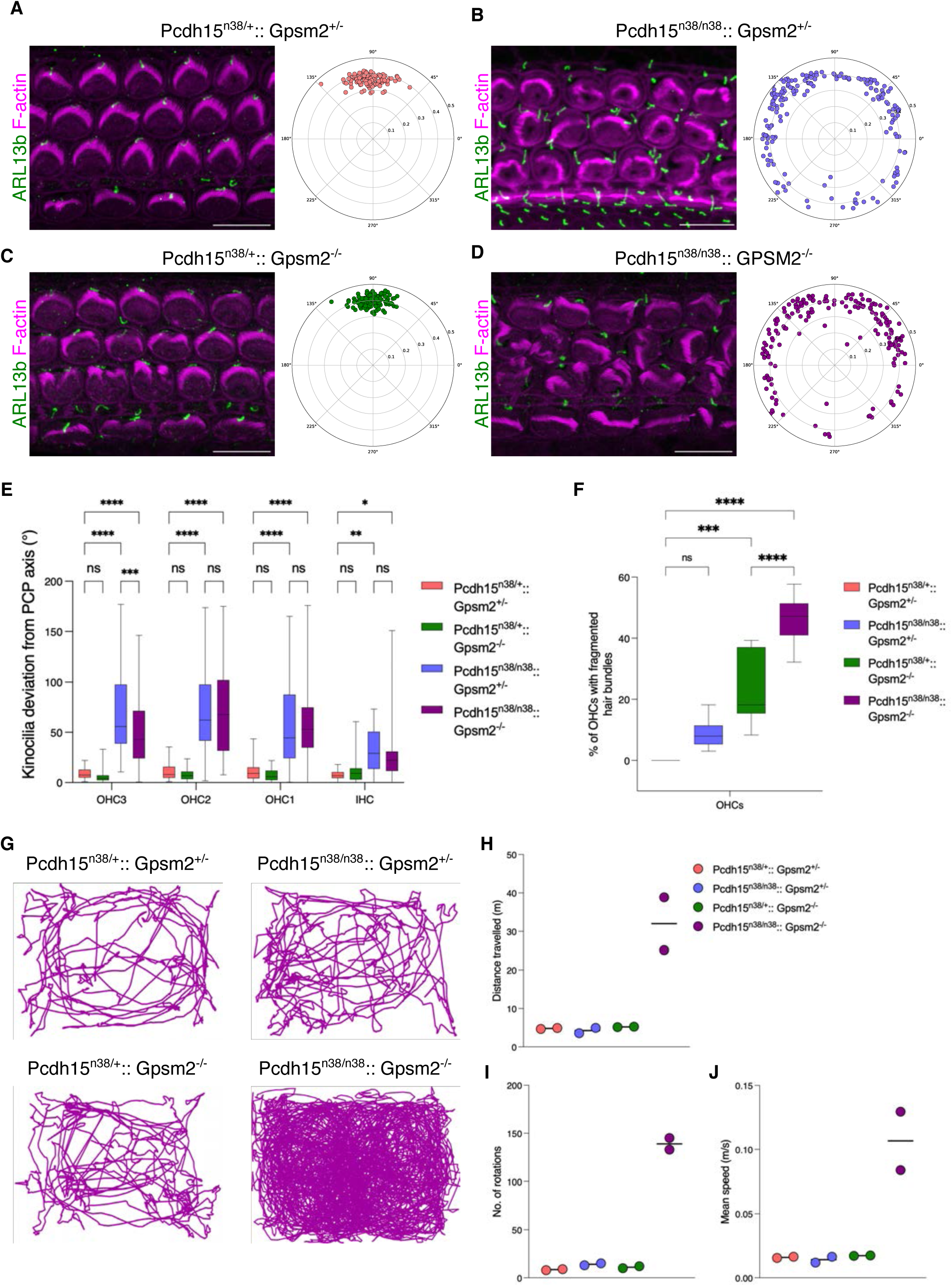
Pcdh15-CD2 and GPSM2 proteins show genetic interaction during hair bundle development. (A-D) Whole mount OC preparation from mice of the indicated genotype at P1 (base). Cilia are marked with ARL13b (green) and stereocilia with phalloidin (magenta) In comparison to Pcdh15^n38/+^:: Gpsm2^+/-^ (A) (pink), the hair bundle polarity of P1 OHCs is perturbed in Pcdh15^n38/n38^:: Gpsm2^+/-^ OC (B) (slate blue) and Pcdh15^n38/n38^:: Gpsm2^-/-^ (D) (dark magenta) OC. (C) Hair bundle polarity is normal in Pcdh15^n38/+^:: Gpsm2^-/-^ (green) OC (Scale bar = 10 μm). (E) Kinocilia are deviated significantly from PCP axis in HCs of Pcdh15^n38/n38^:: Gpsm2^+/-^ (slate blue) & Pcdh15^n38/n38^:: Gpsm2^-/-^ (dark magenta) and aligned along the PCP axis in Pcdh15^n38/+^:: Gpsm2^+/-^ (pink) & Pcdh15^n38/+^:: Gpsm2^-/-^ (green) OC (P1, base). (Two-way Anova with Tukey’s multiple comparisons test, ns = P>0.05, * = P<0.05, ** = P<0.01, *** = P<0.001, **** = P<0.0001). (F) Hair bundles are fragmented in Pcdh15^n38/+^:: Gpsm2^-/-^ (green) OC which is further exacerbated in Pcdh15^n38/n38^:: Gpsm2^-/-^ (dark magenta) (P1, base). (Ordinary one-way Anova with Tukey’s multiple comparisons test, ns = P>0.05, * = P<0.05, ** = P<0.01, *** = P<0.001, **** = P<0.0001). (G) Circling activity is observed only in Pcdh15^n38/n38^:: Gpsm2^-/-^ mice and not in Pcdh15^n38/+^:: Gpsm2^+/-^ or Pcdh15^n38/+^:: Gpsm2^-/-^ mice as compared to control mice when observed under open-field test (OFT) for 5 min. (H, I & J) Pcdh15^n38/n38^:: Gpsm2^-/-^ mice cover more distance, move faster and rotate more than control animals (each circle is an individual animal).

To understand further the nature of this interaction, we investigated the expression of Gpsm2 in Pcdh15 mutants. We find that in HC of Pcdh15^n38/n38^ mice cochlea, GPSM2 expression is unchanged (Fig. 7A). Gpsm2 directs the restricted expression of Gαi which shows a medial expansion in Pcdh15^n38/n38^ (Fig. 5A-C). Gαi interacts with the coiled-coil domain containing proteins, Ccdc88a (Girdin) and Ccdc88c (Daple), we investigated their localisation [35]. We find that the localisation is in the lateral HC junction closest to the kinocilium, and is unchanged in Pcdh15^n38/n38^ cochlea (Fig. 7B). We next asked if the expression of PCDH15 is altered in Gpsm2^-/-^ cochlea. We first used an N-terminal specific antibody that recognises all the major isoforms of PCDH15. We found that expression is unperturbed in Gpsm2^-/-^ HC (Fig. 7C). To ask if the CD2 isoform was affected, we crossed Pcdh15^n38YF/n38YF^ into Gpsm2^-/-^ background and assessed the expression using the HA-tag. We find that PCDH15-CD2-HA still localises to the tip of stereocilia in these mutants (Fig. 7D). These data suggest that although Pcdh15 and Gpsm2 function together, the point of interaction is likely in the downstream pathways of these two molecules.

**Figure 7.**
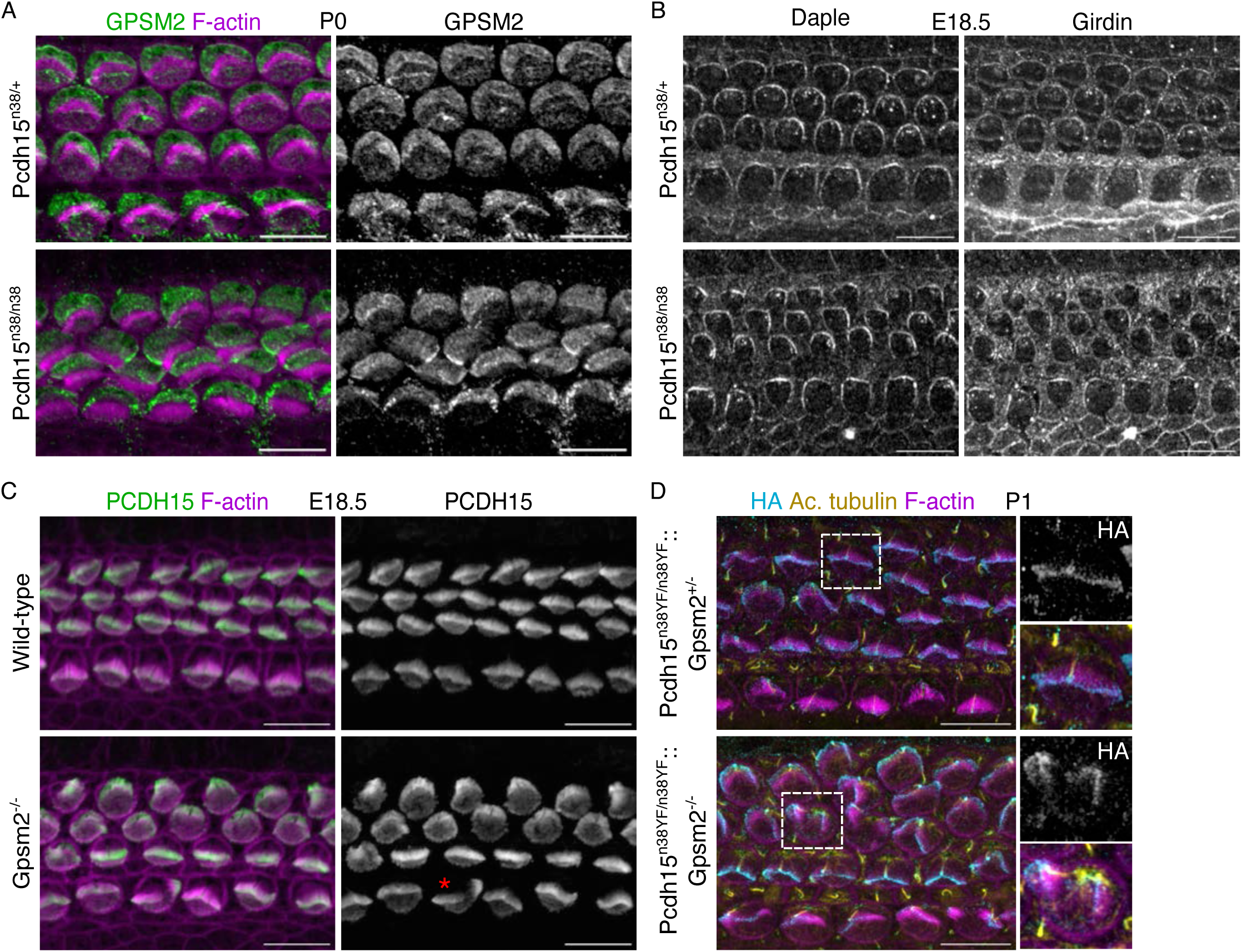
PCDH15-CD2 and GPSM2 proteins operate in parallel pathways during hair bundle development. (A) GPSM2 protein expression (green) complements stereocilia (magenta, marked with phalloidin and is misoriented with hair bundles in P0 (base) HCs of Pcdh15^n38/n38^ mice. (B) Daple & Girdin proteins (grey) are localised on the neural edge of the HCs of both Pcdh15^n38/+^ and Pcdh15^n38/n38^ mice (E18.5, base). (C) PCDH15 protein (green) is localised on the hair bundles (even in fragmented hair bundles, marked with *) of Gpsm2 mutant HCs (E18.5, mid). (D) PCDH15-CD2-HA (cyan) is localised on both kinocilium (yellow, marked with acetylated tubulin) and stereocilia (magenta, marked with phalloidin) in Gpsm2 mutant HCs (P1, base). (Scale bar = 10 μm).

### PCDH15-CD2 expression rescues the intrinsic hair bundle polarity defects

Recently, mini PCDH15-CD2 mediated rescue was reported in a Pcdh15 mutant mouse. Here, the Pcdh15 allele had been removed using a Myo15-cre [36]. Cre is active after intrinsic polarity has already developed in these mice. We asked if re-introducing Pcdh15 function could rescue the intrinsic polarity defects observed in Pcdh15^n38/n38^. We reasoned that by controlling the timing of Cre expression, we could induce recombination such that the mutant Pcdh15^n38/n38^ was replaced by the functional Pcdh15^n38YF/n38YF^ allele (Fig. 2A). We used an Atoh1-CreERT2 line, in which Cre can be activated by tamoxifen induction. We injected dams carrying E17.5 Pcdh15^n38/n38^ mutants with a 5mg/40g dose of tamoxifen and determined the resulting phenotype at E19.5 (Fig. 8A).

**Figure 8.**
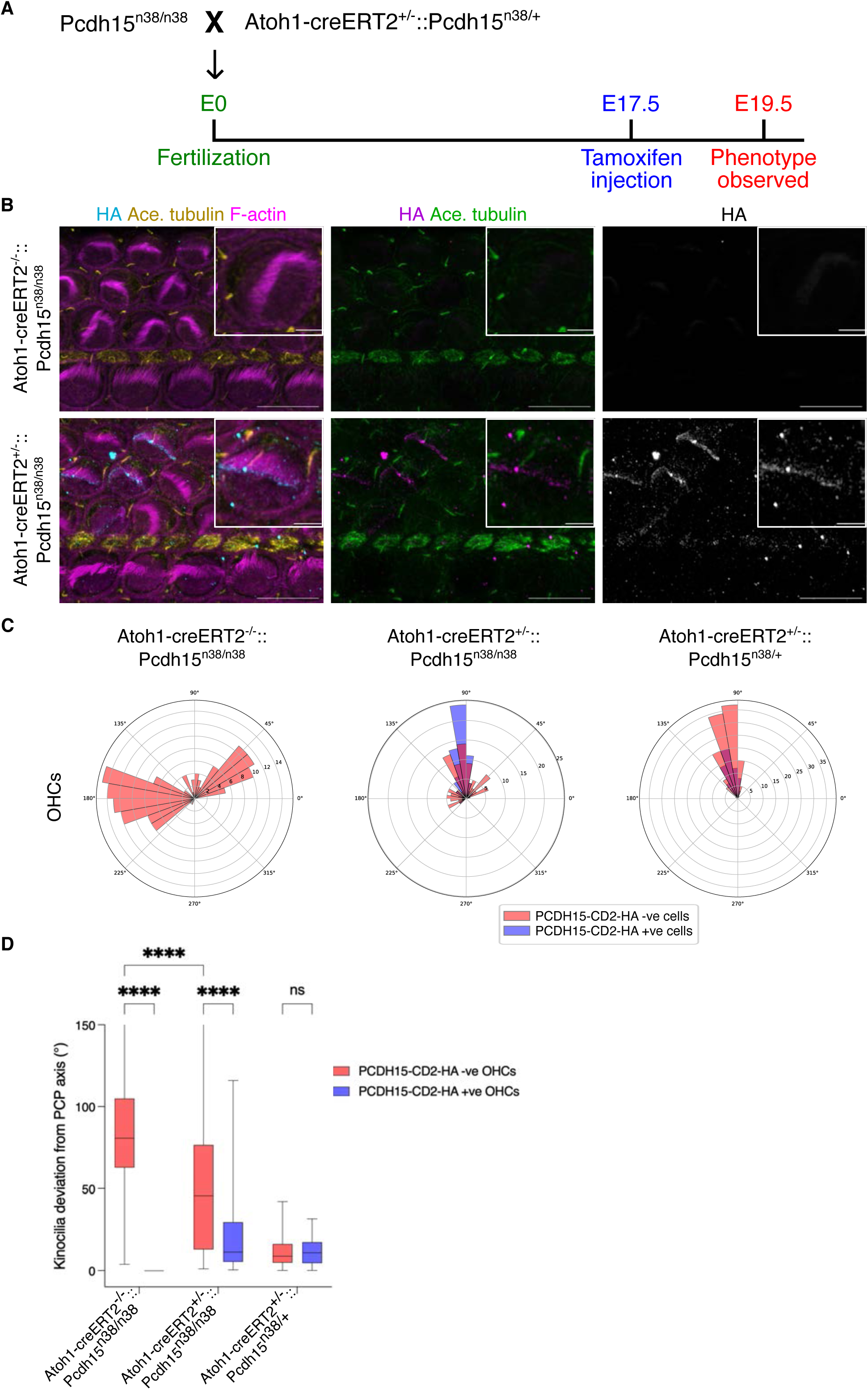
Re-initiation of PCDH15-CD2 expression rescues the intrinsic hair bundle polarity defects. (A) Schematic shows the experimental scheme for the rescue of PCDH15-CD2 expression. (B) PCDH15-CD2-HA (cyan) expression is rescued upon Atoh1-creER induction. Kinocilium (yellow, marked with acetylated tubulin) is associated with stereocilia (magenta, marked with phalloidin), specifically in HCs with PCDH15-CD2-HA expression (E19.5, base). (Scale bar = 10 μm). (C) Circular histogram shows rescue in hair bundle polarity upon Atoh1creER-mediated induction of PCDH15-CD2-HA (slate blue) (E19.5, base). (D) Kinocilia deviation from the PCP axis is improved in HCs with PCDH15-CD2-HA expression (slate blue) (E19.5, base). (Two-way Anova with Tukey’s multiple comparisons test, ns = P>0.05, * = P<0.05, ** = P<0.01, *** = P<0.001, **** = P<0.0001).

In Atoh1-CreET2^-/-^::Pcdh15^n38/n38^ littermate controls, all HC showed kinocilia decoupling from stereocilia, and HC showed intrinsic polarity defects (Fig. 8B-D). These cochlea did not express PCDH15-CD2-HA (Fig. 8B-C). In Atoh1-CreET2^+/-^::Pcdh15^n38/n38^ animals, while Atoh1-CreERT2 induction failed to induce recombination in IHC (Fig. S4A), we observed mosaic expression of PCDH15-CD2-HA in OHC (Fig. 8B-C). In HA positive HC we found that kinocilia and stereocilia were coupled (Fig. 8B). We quantified the kinocilium position in hair cells, differentiating whether they expressed PCDH15-CD2-HA. We found that the kinocilia of PCDH15-CD2-HA expressing OHC align with the tissue axis (26° ± 30°), in contrast to the non-expressing OHC (48° ± 37°) (Fig. 8D).

## DISCUSSION

The initial motivation of our study was to investigate the role of PCDH15-CD2 phosphorylation in the development and physiology of hair cells. To determine this we had engineered a mouse in which epitope-tagged CD-2 and phospho-mutant CD-2 domains could be conditionally expressed. However in this mouse, the unmutated PCDH15-CD2 (n38) could not be expressed. Moreover, our analysis revealed that tyrosine phosphorylation of the CD2 domain likely does not play a role in Pcdh15 function. However, using the null n38, and the functional and tagged n38YF alleles, we could analyse the early mutant phenotype in greater resolution and obtain insights into the earliest expression of PCDH15-CD2.

As has been described previously, mutants of Pcdh15-CD2 show early intrinsic polarity defects [21, 22]. By E16.5, Pcdh15-n38 show more variance than either heterozygotes or Pcdh15-n38YF. This has been thought to be due to the absence of kinocilial links, a conjecture supported by mutations in the partner of Pcdh15, Cdh23 [22]. These also show intrinsic polarity defects. However, kinocilial links appear later in development, and are only first visible in the IHC found at the base of the cochlea at E17.5 [25]. This is after the initial centrifugal relocalisation of the kinocilia [8–10]. This may imply that Pcdh15 contributes to the generation or maintenance of intrinsic planar polarity, independent of its role in the kinocilial links. Using the epitope tagged allele that we had generated, we could detect the earliest localisation of PCDH15-CD2. Initial expression was found as early as E15.5 (IHCs) before the off-centre movement of the kinocilium, in and around the base of the cilium. Similar expression was also observed for OHC and for HCs of the vestibular organs.

During the continued development of hair cells, the kinocilium shifts along the periphery such that it is in the centre of the crescent made by Gαi3 expression [9, 10]. This occurs between E16.5 to E18.5. In Pcdh15-n38 mutants, this shift does not occur. This does indicate a coupling between the Gαi3 domain and the kinocilium, but the nature of this interaction is unclear. A similar failure in Gαi3 and kinocilia correlation has been observed in two other mutants, the basal-body protein Mkks, and an interactor of Gαi3, Daple [10, 35]. Daple, and a related Gαi3 interacting protein, Girdin, do not show significant alteration in expression in Pcdh15-n38. However, Gαi3 expression in Pcdh15n38 spreads medially. This has also been observed in Mkks mutants and in other ciliary mutants, such as Ift88 and Bbs8 [10, 14, 17]<u>. (BbS8 – Vangl2)</u> In addition to Mkks, Ift88 and Bbs8, a number of mutants in kinociliary genes, including Ift20, Ift27, and Kif3a show variation in stereocilia bundle orientation, to differing extents [17, 18]. Kinocilia position can influence planar polarity. Bbs8 is known to influence the trafficking of Vangl2, and overall epithelial organisation may be affected as a result [17]. In our analysis of Pcdh15-n38 mutants, we did not detect an alteration in Vangl2 localisation. Recent data suggests that the kinocilia’s position alters junctional mechanics during chick cochlea extension, altering polarity [37]. Pcdh15-CD2 may alter the ability of the kinocilia to influence neighbouring cellular dynamics leading to defects in planar polarity.

Of note is the analysis of the basal body position performed in the Kif3a mutant. In wild-type hair cells, the basal body of the kinocilia is apical, and the daughter centriole more basal. In Kif3a mutants, that difference in apicobasal position is lost [18]. In Pcdh15-CD2 mutants, the centrioles of the basal body are mispositioned. The mispositioning of the basal body/pericentriolar matrix may impact the restriction of Gαi3, and may also impact the peripheral shift of the kinocilia so that it is coincident with the centre of the Gαi3 crescent. Perturbed Rac-PAK signalling is thought to underlie the perturbed basal body position in the Kif3a mutant, with active Rac1 recruited to the pericentriolar matrix through the microtubule-associated protein, Lis1 [38]. Lis1 is also required for the dynein localisation around the kinocilium centrosome. Our unpublished data suggests that PCDH15-CD2 can interact directly with dynein in vitro.

Generating the apical blueprint of the hair cell requires coordination between the kinocilium and the developing stereociliary bundles [11]. Ciliary genes and the PCDH15-CD2 isoform direct kinocilium development, whereas the GPSM2/Gαi proteins specify the generation of the stereociliary bundle. How these two systems are coupled is, however, unknown. Despite the localisation of the two molecules being independent, compound homozygotes of Pcdh15n38 and Gpsm2 genes showed a genetic interaction. Although kinocilia position is similarly perturbed in Pcdh15-CD2 and double mutants, hair bundle fragmentation is exacerbated in the double mutants. An observation from studies on Daple mutants suggests that fragmentation occurs when the initial centrifugal migration of the kinocilia is to a point that is outside of the Gαi3 crescent [35]. It is possible that this may also underlie the increased fragmentation we observe in the Pcdh15n38/Gpsm2 double mutants. Both single and double mutants lack a Preyer’s reflex, indicative of hearing loss. Individually, Pcdh15-n38 and Gpsm2 null do not show defects in vestibular behaviours. However, double mutants showed hyperactivity and circling. The reason for circling is not immediately apparent, as we were unable to detect differences in vestibular hair cells. This is in contrast to earlier analysis of the mPins allele of Gpsm2 [39]. Here, a vestibular phenotype is observed in the single mutant, and utricular hair cells show reduced stereocilia length. The reason for the discrepancy between the two alleles is unclear. However, the mPins allele may produce a truncated protein, which could act as a dominant negative.

Using Cre mediated recombination, PCDH15-CD2 expression can be restored. Using this feature, the alignment of polarity and kinocilia-stereocilia coupling could be rescued in hair cells in which PCDH15-CD2-HA expression was induced. Due to the low efficacy of the Atoh-CreER2 at post-natal stages, we could only rescue intrinsic polarity until E19.5. However, this does suggest that PCDH15-CD2 plays a central role in the coordination of the kinocilia and stereociliary bundles and in the generation of intrinsic polarity through a mechanism likely independent of initial kinocilia assembly and likely independent of the formation of the kinocilial links.

## Supporting information

Supp Move 1-4

Supp Move 1-4

Supp Move 1-4

Supp Move 1-4

## ACKNOWLEDGMENTS

This work was supported by the Department of Atomic Energy, Government of India, Project Identification No. RTI 4006, and grants from the Royal National Institute for Deaf People (IPG programme), SERB and TIFR Infosys-Leading Edge Grant. We acknowledge the support of the Animal Care and Resources Centre, Electron Microscopy Facility and Central Imaging Facility at NCBS. A.P. acknowledges Simons Foundation International for support through Simons-Ashoka early career fellowship. We thank members of the Ear Lab at NCBS for their feedback and discussions.

## SUPPLEMENTARY FIGURE LEGENDS

**Figure S1.**
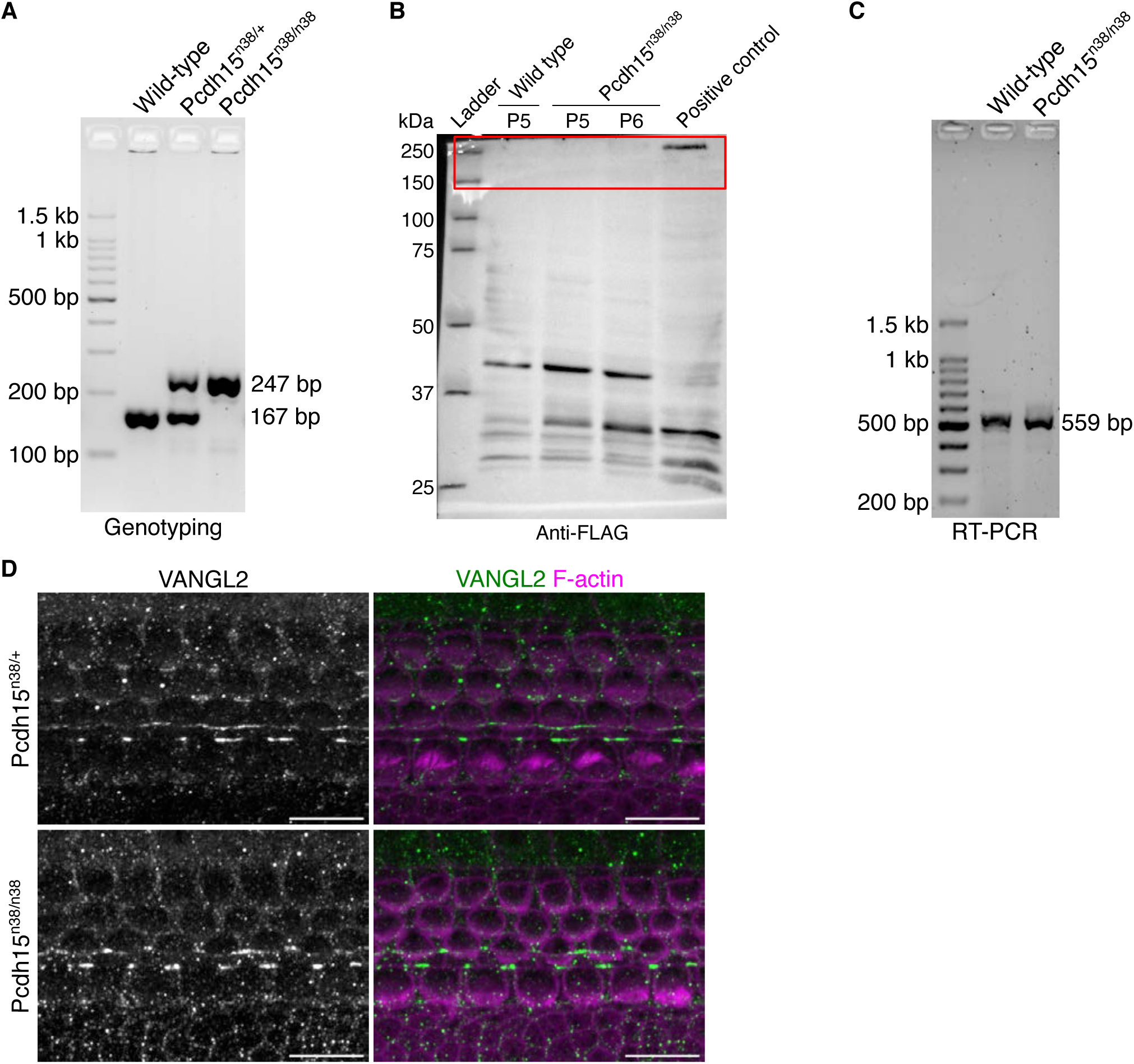
Pcdh15^n38/n38^ mice are functionally null and act similar to Pcdh15-ΔCD2 mutants. (A) A small fragment of Pcdh15 exon 38 was amplified in WT (167 bp) and Pcdh15^n38/n38^ (247 bp) animals for genotyping. (B) Full-length western blot of the Fig. 1C. (C) mRNA level for CD2 domain is normal in Pcdh15^n38/n38^ cochlea. (D) VANGL2 protein (green) localisation is unperturbed in Pcdh15^n38/n38^ OC (E17.5, base). (Scale bar = 10 μm).

**Figure S2.**
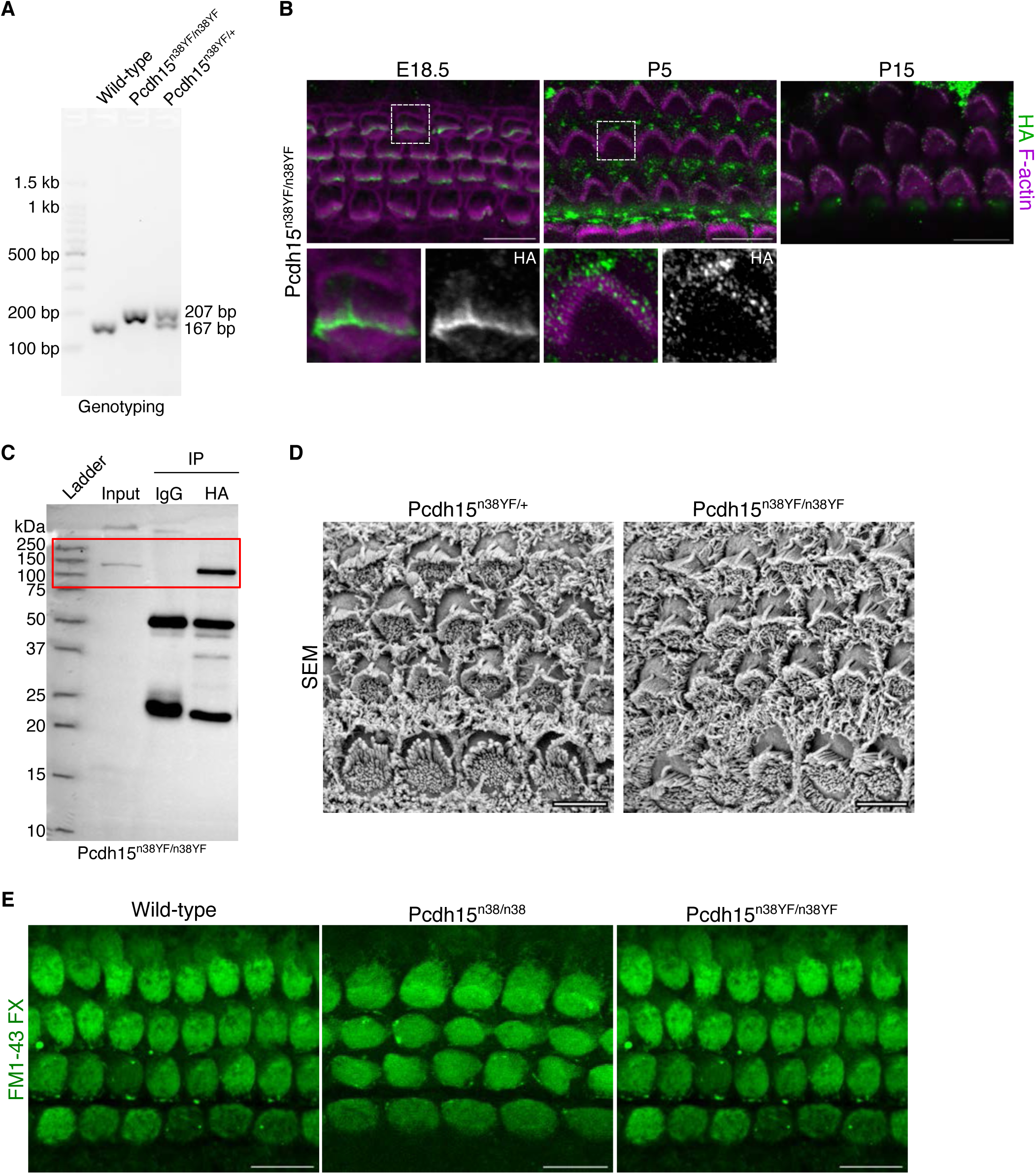
Tyrosine phosphorylation of the CD2 domain is not essential for PCDH15 function. (A) Part of WT and Pcdh15^n38YF/n38YF^ exon 38 are amplified in different sizes (167 bp and 207 bp respectively) with same primer pair for genotyping. (B) PCDH15-CD2-HA protein (green) is localised at the tip of stereocilia (magenta, marked with phalloidin) at E18.5 (mid). At P5 (mid), this expression is localised in all three rows of stereocilia and refined more in P15 stereocilia (mid). (Scale bar = 10 μm). (C) Full-length western blot of the Fig. 2E. (D) SEM imaging shows normal hair bundle polarity in both Pcdh15^n38/+^ and Pcdh15^n38YF/n38YF^ HCs (P0, mid). (Scale bar = 5 μm). (E) MET channels are open in HCs of Pcdh15^n38/n38^ and Pcdh15^n38YF/n38YF^ mice at P5. FM1-43 FX dye (green) is used to detect open MET channels. (Scale bar = 10 μm).

**Figure S3.**
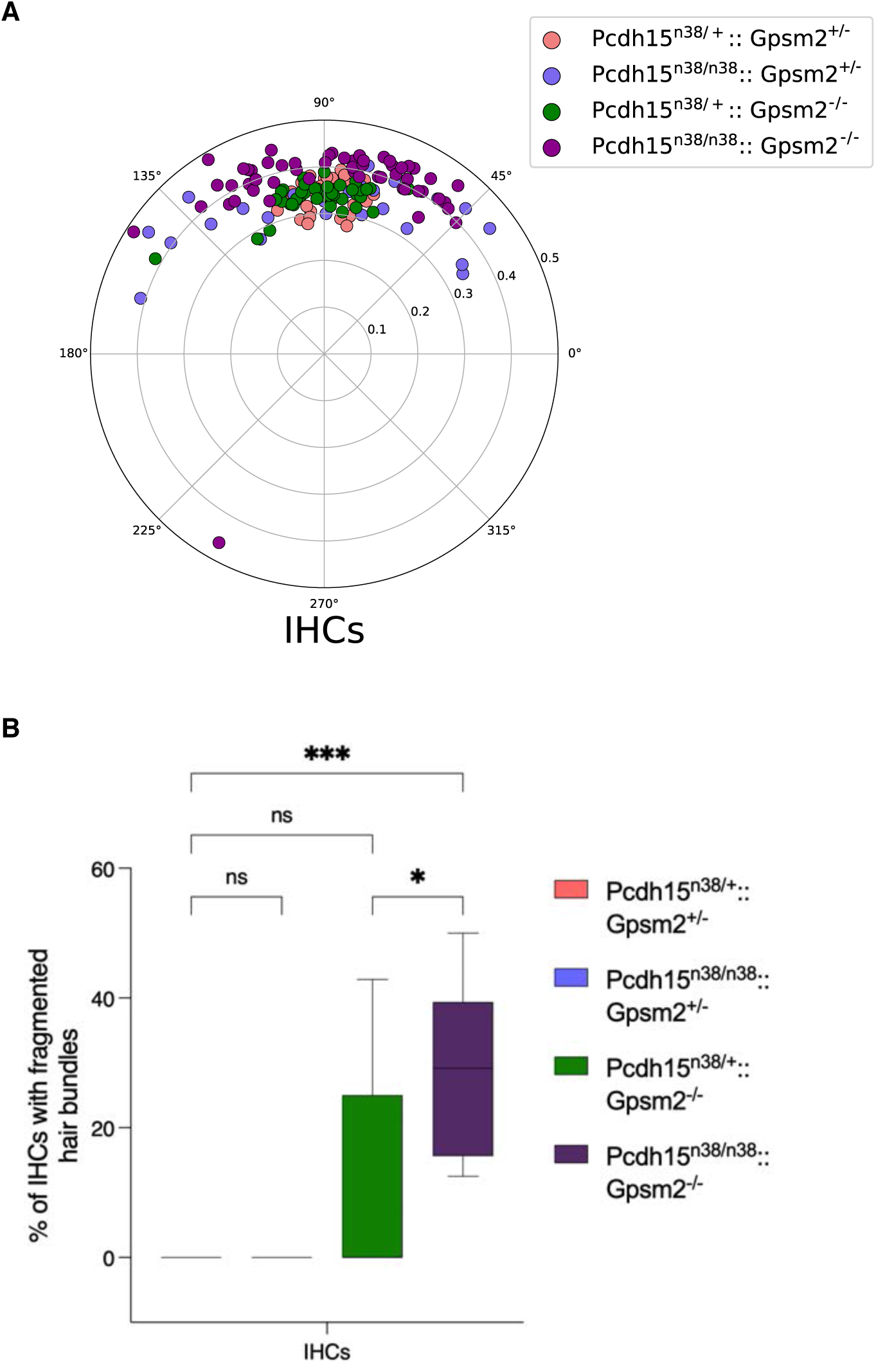
Pcdh15-CD2 and Gpsm2 show genetic interaction during hair bundle development. (A) In comparison to Pcdh15^n38/+^:: Gpsm2^+/-^ (pink), the hair bundle polarity (polar projections) is perturbed in P1 IHCs of Pcdh15^n38/n38^:: Gpsm2^+/-^ (slate blue) and Pcdh15^n38/n38^:: Gpsm2^-/-^ (dark magenta) OC (base). Hair bundle polarity is normal in Pcdh15^n38/+^:: Gpsm2^-/-^ (green) OC. (B) Hair bundles are fragmented in IHCs of only Pcdh15^n38/n38^:: Gpsm2^-/-^ (dark magenta). (Ordinary one-way Anova with Tukey’s multiple comparisons test, ns = P>0.05, * = P<0.05, ** = P<0.01, *** = P<0.001, **** = P<0.0001).

## SUPPLEMENTARY TABLES

**Table S1.**
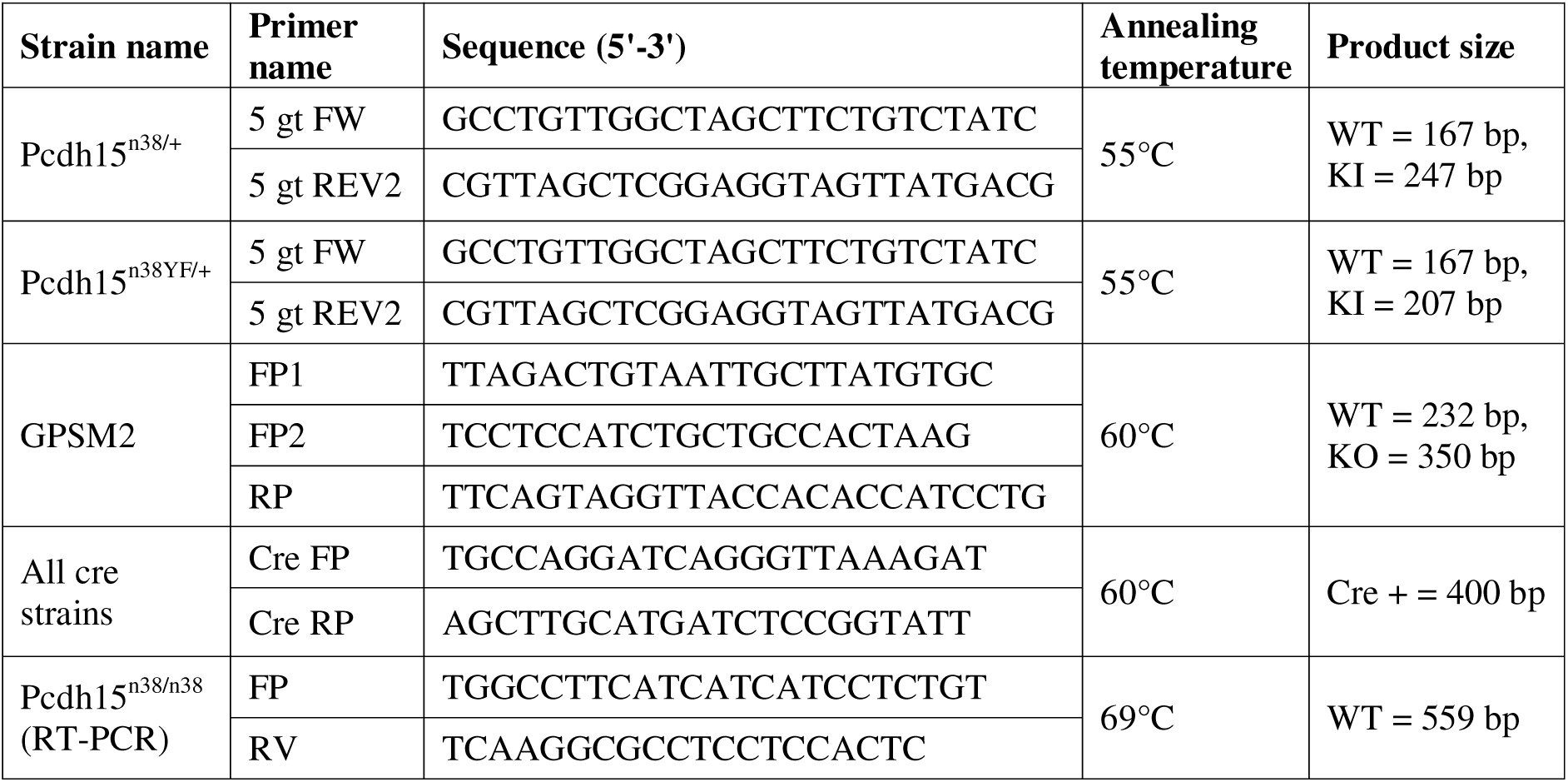
Genotyping and RT-PCR conditions for different genes used. These include their specific primers, annealing temperature, and amplified product size.

**Table S2.**
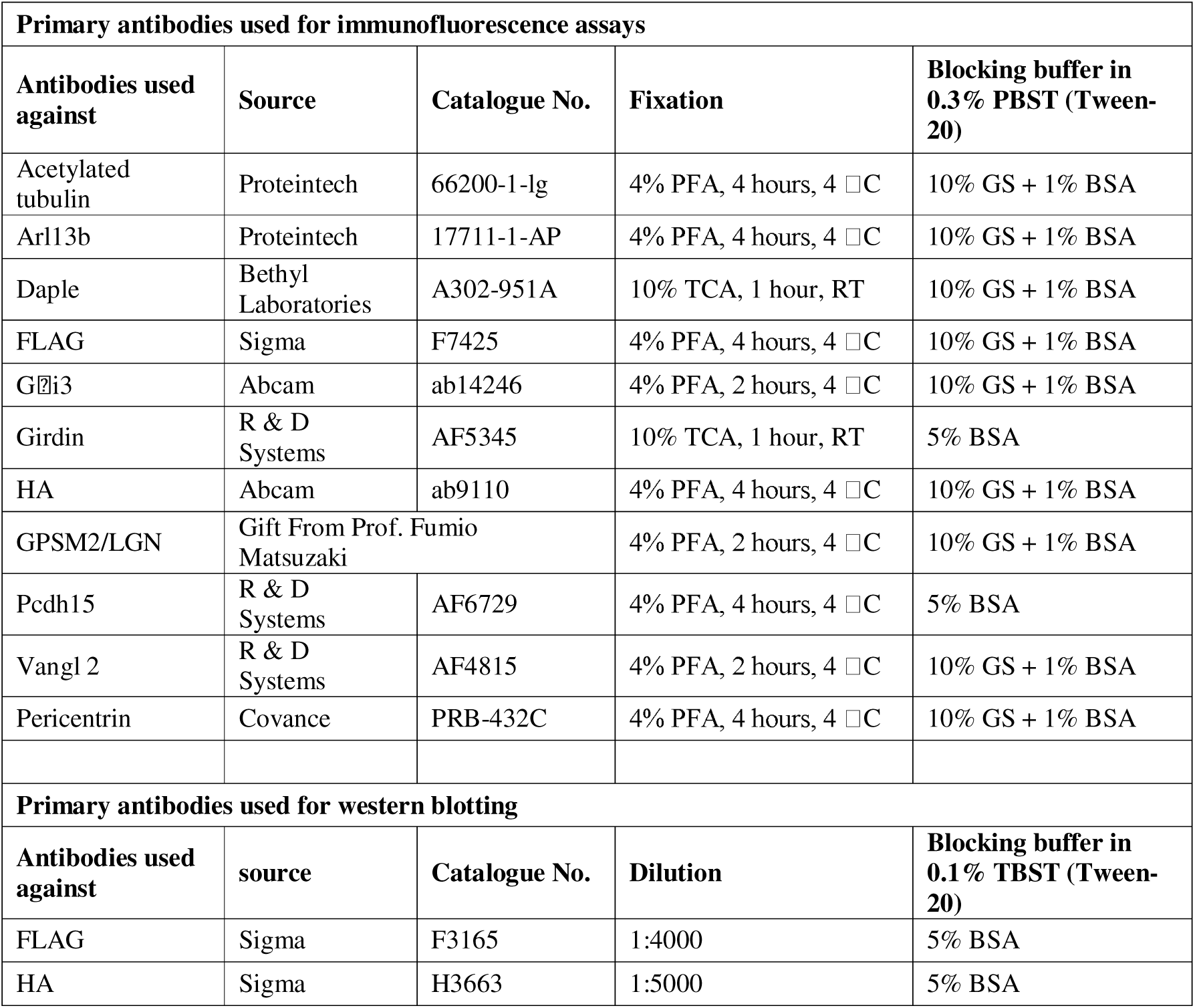
List of primary antibodies used for immunofluorescence assays and western blotting.

**Table S3.** List of secondary antibodies used for immunofluorescence assays.

| **Antibodies** | **source** | **Catalogue No.** |
| --- | --- | --- |
| Alexa Fluor™ 488 Phalloidin | Thermo Fisher Scientific | A-12379 |
| Alexa Fluor™ 568 Phalloidin | Thermo Fisher Scientific | A-12380 |
| Donkey anti-Mouse IgG Alexa Fluor™ 488 | Thermo Fisher Scientific | A-21202 |
| Donkey anti-Mouse IgG Alexa Fluor™ 594 | Thermo Fisher Scientific | A-21245 |
| Donkey anti-Sheep IgG Secondary Antibody, Alexa Fluor™ 488 | Thermo Fisher Scientific | A-11015 |
| Goat anti-chick IgG Secondary Antibody, Alexa Fluor™ 594 | Thermo Fisher Scientific | A-11042 |
| Goat anti-Mouse IgG Secondary Antibody, Alexa Fluor™ 488 | Thermo Fisher Scientific | A-11001 |
| Goat anti-Mouse IgG Secondary Antibody, Alexa Fluor™ 594 | Thermo Fisher Scientific | A-11032 |
| Goat anti-Mouse IgG Secondary Antibody, Alexa Fluor™ 647 | Thermo Fisher Scientific | A-21235 |
| Goat anti-Rabbit IgG Secondary Antibody, Alexa Fluor™ 488 | Thermo Fisher Scientific | A-11008 |
| Goat anti-Rabbit IgG Secondary Antibody, Alexa Fluor™ 647 | Thermo Fisher Scientific | A-21245 |
| Goat anti-Rat IgG Secondary Antibody, Alexa Fluor™ 488 | Thermo Fisher Scientific | A-11007 |

## REFERENCES

1. Dror AA, Avraham KB. Hearing loss: mechanisms revealed by genetics and cell biology. Annual review of genetics. 2009;43:411–37. doi: 10.1146/annurev-genet-102108-134135. PubMed PMID: 19694516.

2. Driver EC, Kelley MW. Development of the cochlea. Development. 2020;147(12). Epub 20200622. doi: 10.1242/dev.162263. PubMed PMID: 32571852; PubMed Central PMCID: PMCPMC7327997.

3. Tilney LG, Tilney MS, DeRosier DJ. Actin filaments, stereocilia, and hair cells: how cells count and measure. Annu Rev Cell Biol. 1992;8:257–74. Epub 1992/01/01. doi: 10.1146/annurev.cb.08.110192.001353. PubMed PMID: 1476800.

4. Shotwell SL, Jacobs R, Hudspeth AJ. Directional sensitivity of individual vertebrate hair cells to controlled deflection of their hair bundles. Ann N Y Acad Sci. 1981;374:1–10. doi: 10.1111/j.1749-6632.1981.tb30854.x. PubMed PMID: 6978627.

5. Tilney LG, Tilney MS, Saunders JS, DeRosier DJ. Actin filaments, stereocilia, and hair cells of the bird cochlea. III. The development and differentiation of hair cells and stereocilia. Dev Biol. 1986;116(1):100–18. Epub 1986/07/01. doi: 0012-1606(86)90047-3 [pii]. PubMed PMID: 3732601.

6. Kaltenbach JA, Falzarano PR, Simpson TH. Postnatal development of the hamster cochlea. II. Growth and differentiation of stereocilia bundles. J Comp Neurol. 1994;350(2):187–98. doi: 10.1002/cne.903500204. PubMed PMID: 7884037.

7. Zine A, Romand R. Development of the auditory receptors of the rat: a SEM study. Brain Res. 1996;721(1-2):49–58. doi: 10.1016/0006-8993(96)00147-3. PubMed PMID: 8793083.

8. Chakravarthy SR, van Zanten TS, Ladher RK. Initiation and Formation of Stereocilia during the Development of Mouse Cochlear Hair Cells. bioRxiv. 2024:2024.03.23.586377. doi: 10.1101/2024.03.23.586377.

9. Tarchini B, Jolicoeur C, Cayouette M. A molecular blueprint at the apical surface establishes planar asymmetry in cochlear hair cells. Dev Cell. 2013;27(1):88–102. doi: 10.1016/j.devcel.2013.09.011. PubMed PMID: 24135232.

10. Ezan J, Lasvaux L, Gezer A, Novakovic A, May-Simera H, Belotti E, et al. Primary cilium migration depends on G-protein signalling control of subapical cytoskeleton. Nat Cell Biol. 2013;15(9):1107–15. doi: 10.1038/ncb2819. PubMed PMID: 23934215.

11. Tarchini B, Lu X. New insights into regulation and function of planar polarity in the inner ear. Neurosci Lett. 2019;709:134373. Epub 20190708. doi: 10.1016/j.neulet.2019.134373. PubMed PMID: 31295539; PubMed Central PMCID: PMCPMC6732021.

12. Montcouquiol M, Kelley MW. Development and Patterning of the Cochlea: From Convergent Extension to Planar Polarity. Cold Spring Harb Perspect Med. 2020;10(1). Epub 20200102. doi: 10.1101/cshperspect.a033266. PubMed PMID: 30617059; PubMed Central PMCID: PMCPMC6938661.

13. Morin X, Bellaiche Y. Mitotic spindle orientation in asymmetric and symmetric cell divisions during animal development. Dev Cell. 2011;21(1):102–19. doi: 10.1016/j.devcel.2011.06.012. PubMed PMID: 21763612.

14. Bhonker Y, Abu-Rayyan A, Ushakov K, Amir-Zilberstein L, Shivatzki S, Yizhar-Barnea O, et al. The GPSM2/LGN GoLoco motifs are essential for hearing. Mammalian genome : official journal of the International Mammalian Genome Society. 2016;27(1-2):29–46. Epub 20151211. doi: 10.1007/s00335-015-9614-7. PubMed PMID: 26662512; PubMed Central PMCID: PMCPMC5913375.

15. Ross AJ, May-Simera H, Eichers ER, Kai M, Hill J, Jagger DJ, et al. Disruption of Bardet-Biedl syndrome ciliary proteins perturbs planar cell polarity in vertebrates. Nat Genet. 2005;37(10):1135–40. Epub 2005/09/20. doi: ng1644 [pii] 10.1038/ng1644. PubMed PMID: 16170314.

16. Jones C, Roper VC, Foucher I, Qian D, Banizs B, Petit C, et al. Ciliary proteins link basal body polarization to planar cell polarity regulation. Nat Genet. 2008;40(1):69–77. Epub 2007/12/11. doi: ng.2007.54 [pii] 10.1038/ng.2007.54. PubMed PMID: 18066062.

17. May-Simera HL, Petralia RS, Montcouquiol M, Wang YX, Szarama KB, Liu Y, et al. Ciliary proteins Bbs8 and Ift20 promote planar cell polarity in the cochlea. Development. 2015;142(3):555–66. doi: 10.1242/dev.113696. PubMed PMID: 25605782; PubMed Central PMCID: PMCPMC4302998.

18. Sipe CW, Lu X. Kif3a regulates planar polarization of auditory hair cells through both ciliary and non-ciliary mechanisms. Development. 2011;138(16):3441–9. Epub 20110713. doi: 10.1242/dev.065961. PubMed PMID: 21752934; PubMed Central PMCID: PMCPMC3143564.

19. Jagger D, Collin G, Kelly J, Towers E, Nevill G, Longo-Guess C, et al. Alstrom Syndrome protein ALMS1 localizes to basal bodies of cochlear hair cells and regulates cilium-dependent planar cell polarity. Hum Mol Genet. 2011;20(3):466–81. Epub 20101111. doi: 10.1093/hmg/ddq493. PubMed PMID: 21071598; PubMed Central PMCID: PMCPMC3016908.

20. Richardson GP, Petit C. Hair-Bundle Links: Genetics as the Gateway to Function. Cold Spring Harb Perspect Med. 2019;9(12). Epub 20191202. doi: 10.1101/cshperspect.a033142. PubMed PMID: 30617060; PubMed Central PMCID: PMCPMC6886453.

21. Pawlowski KS, Kikkawa YS, Wright CG, Alagramam KN. Progression of inner ear pathology in Ames waltzer mice and the role of protocadherin 15 in hair cell development. J Assoc Res Otolaryngol. 2006;7(2):83–94. doi: 10.1007/s10162-005-0024-5. PubMed PMID: 16408167; PubMed Central PMCID: PMCPMC2504581.

22. Lefevre G, Michel V, Weil D, Lepelletier L, Bizard E, Wolfrum U, et al. A core cochlear phenotype in USH1 mouse mutants implicates fibrous links of the hair bundle in its cohesion, orientation and differential growth. Development. 2008;135(8):1427–37. Epub 20080313. doi: 10.1242/dev.012922. PubMed PMID: 18339676.

23. Webb SW, Grillet N, Andrade LR, Xiong W, Swarthout L, Della Santina CC, et al. Regulation of PCDH15 function in mechanosensory hair cells by alternative splicing of the cytoplasmic domain. Development. 2011;138(8):1607–17. Epub 2011/03/24. doi: 138/8/1607 [pii] 10.1242/dev.060061. PubMed PMID: 21427143.

24. Honda A, Kita T, Seshadri SV, Misaki K, Ahmed Z, Ladbury JE, et al. FGFR1-mediated protocadherin-15 loading mediates cargo specificity during intraflagellar transport in inner ear hair-cell kinocilia. Proc Natl Acad Sci U S A. 2018;115(33):8388–93. Epub 20180730. doi: 10.1073/pnas.1719861115. PubMed PMID: 30061390; PubMed Central PMCID: PMCPMC6099903.

25. Goodyear RJ, Marcotti W, Kros CJ, Richardson GP. Development and properties of stereociliary link types in hair cells of the mouse cochlea. J Comp Neurol. 2005;485(1):75–85. doi: 10.1002/cne.20513. PubMed PMID: 15776440.

26. Abe T, Inoue KI, Furuta Y, Kiyonari H. Pronuclear Microinjection during S-Phase Increases the Efficiency of CRISPR-Cas9-Assisted Knockin of Large DNA Donors in Mouse Zygotes. Cell reports. 2020;31(7):107653. doi: 10.1016/j.celrep.2020.107653. PubMed PMID: 32433962.

27. Naito Y, Hino K, Bono H, Ui-Tei K. CRISPRdirect: software for designing CRISPR/Cas guide RNA with reduced off-target sites. Bioinformatics. 2015;31(7):1120–3. Epub 20141120. doi: 10.1093/bioinformatics/btu743. PubMed PMID: 25414360; PubMed Central PMCID: PMCPMC4382898.

28. Lallemand Y, Luria V, Haffner-Krausz R, Lonai P. Maternally expressed PGK-Cre transgene as a tool for early and uniform activation of the Cre site-specific recombinase. Transgenic research. 1998;7(2):105–12. doi: 10.1023/a:1008868325009. PubMed PMID: 9608738.

29. Fujita I, Shitamukai A, Kusumoto F, Mase S, Suetsugu T, Omori A, et al. Endfoot regeneration restricts radial glial state and prevents translocation into the outer subventricular zone in early mammalian brain development. Nat Cell Biol. 2020;22(1):26–37. Epub 20191223. doi: 10.1038/s41556-019-0436-9. PubMed PMID: 31871317.

30. Machold R, Fishell G. Math1 is expressed in temporally discrete pools of cerebellar rhombic-lip neural progenitors. Neuron. 2005;48(1):17–24. doi: 10.1016/j.neuron.2005.08.028. PubMed PMID: 16202705.

31. Honda A, Freeman SD, Sai X, Ladher RK, O’Neill P. From placode to labyrinth: culture of the chicken inner ear. Methods. 2014;66(3):447–53. Epub 2013/06/25. doi: 10.1016/j.ymeth.2013.06.011. PubMed PMID: 23792918.

32. Singh N, Prakash A, Chakravarthy SR, Kaushik R, Ladher RK. In Ovo and Ex Ovo Methods to Study Avian Inner Ear Development. Journal of visualized experiments : JoVE. 2022;(184). Epub 20220616. doi: 10.3791/64172. PubMed PMID: 35786636.

33. Pepermans E, Michel V, Goodyear R, Bonnet C, Abdi S, Dupont T, et al. The CD2 isoform of protocadherin-15 is an essential component of the tip-link complex in mature auditory hair cells. EMBO Mol Med. 2014;6(7):984–92. doi: 10.15252/emmm.201403976. PubMed PMID: 24940003; PubMed Central PMCID: PMCPMC4119359.

34. Gale JE, Marcotti W, Kennedy HJ, Kros CJ, Richardson GP. FM1-43 dye behaves as a permeant blocker of the hair-cell mechanotransducer channel. J Neurosci. 2001;21(18):7013–25. PubMed PMID: 11549711.

35. Siletti K, Tarchini B, Hudspeth AJ. Daple coordinates organ-wide and cell-intrinsic polarity to pattern inner-ear hair bundles. Proc Natl Acad Sci U S A. 2017;114(52):E11170–E9. Epub 20171211. doi: 10.1073/pnas.1716522115. PubMed PMID: 29229865; PubMed Central PMCID: PMCPMC5748220.

36. Ivanchenko MV, Hathaway DM, Mulhall EM, Booth KT, Wang M, Peters CW, et al. PCDH15 Dual-AAV Gene Therapy for Deafness and Blindness in Usher Syndrome Type 1F Models. The Journal of clinical investigation. 2024;134(23). Epub 20241015. doi: 10.1172/JCI177700. PubMed PMID: 39441757; PubMed Central PMCID: PMCPMC11601915.

37. Prakash A, Weninger J, Singh N, Raman S, Kruse K, Rao M, et al. Junctional Force Patterning drives both Positional and Orientational Order in Auditory Epithelia. Research Square. 2023. doi: 10.21203/rs.3.rs-2508957/v1.

38. Sipe CW, Liu L, Lee J, Grimsley-Myers C, Lu X. Lis1 mediates planar polarity of auditory hair cells through regulation of microtubule organization. Development. 2013;140(8):1785–95. doi: 10.1242/dev.089763. PubMed PMID: 23533177; PubMed Central PMCID: PMCPMC3621493.

39. Mauriac SA, Hien YE, Bird JE, Carvalho SD, Peyroutou R, Lee SC, et al. Defective Gpsm2/Galpha(i3) signalling disrupts stereocilia development and growth cone actin dynamics in Chudley-McCullough syndrome. Nature communications. 2017;8:14907. Epub 20170407. doi: 10.1038/ncomms14907. PubMed PMID: 28387217; PubMed Central PMCID: PMCPMC5385604.

